# Inflammatory ER Stress Responses Dictate the Immunopathogenic Progression of Systemic Candidiasis

**DOI:** 10.1101/2022.11.17.513879

**Authors:** Deepika Awasthi, Sahil Chopra, Byuri A. Cho, Alexander Emmanuelli, Tito A. Sandoval, Sung-Min Hwang, Chang-Suk Chae, Camilla Salvagno, Chen Tan, Liliana Vasquez-Urbina, Jose J. Fernandez Rodriguez, Sara F. Santagostino, Takao Iwawaki, E. Alfonso Romero-Sandoval, Mariano Sanchez Crespo, Diana K. Morales, Iliyan D. Iliev, Tobias M. Hohl, Juan R. Cubillos-Ruiz

## Abstract

Recognition of pathogen-associated molecular patterns can trigger the IRE1α arm of the endoplasmic reticulum (ER) stress response in immune cells. IRE1α activation has been shown to maintain ER homeostasis while simultaneously coordinating diverse immunomodulatory programs in the setting of bacterial and viral infections. However, the role of IRE1α signaling in innate immune responses to fungal pathogens is unknown. Here we report that systemic infection with the fungus *Candida albicans* causes severe renal immunopathology by triggering inflammatory IRE1α hyperactivation in host myeloid cells. Mechanistically, sensing of fungal β-glucans by the C-type lectin receptor Dectin-1 induced Src–Syk–NOX-dependent accumulation of intracellular reactive oxygen species and the ensuing generation of lipid peroxidation byproducts that sustained IRE1α activation. Selective deletion of IRE1α in leukocytes, or treatment with an IRE1α pharmacological inhibitor, reduced detrimental inflammatory responses in the kidney and extended survival in mice systemically infected with *C. albicans*. Hence, controlling IRE1α overactivation may be useful to impede the fatal immunopathogenic progression of disseminated candidiasis.

**One sentence summary:** Innate IRE1α signaling in disseminated candidiasis

## INTRODUCTION

Sensing and responding to pathogens entails massive protein synthesis, folding, modification, and trafficking in immune cells, which are processes governed by the endoplasmic reticulum (ER). The excess demand in protein handling can lead to accumulation of misfolded proteins in this organelle (*1*). Pathogens can also produce factors that alter the protein-folding capacity of the ER (*2*). Both events can provoke “ER stress”, a cellular state that activates the unfolded protein response (UPR) (*1*). Three ER stress sensors govern the UPR: inositol-requiring enzyme 1 alpha (IRE1α), protein kinase R-like endoplasmic reticulum kinase (PERK) and activating transcription factor 6 (ATF6). IRE1α is the most conserved ER stress sensor (*1*). When ER homeostasis is altered, IRE1α undergoes activation, prompting excision of a 26-nucleotide fragment from the *XBP1* mRNA. The spliced isoform generated (*XBP1s*) codes for the functionally active transcription factor XBP1, which activates various mechanisms to restore ER proteostasis (*1*). The canonical function of IRE1α-XBP1 in the UPR is to promote rapid expression of protein chaperones, foldases, glycosylases, quality control proteins, and ER-associated degradation components to restore ER proteostasis (*1*). However, concomitant with these adaptive responses, IRE1α signaling can also govern a variety of UPR-independent processes and pathways implicated in multiple pathological conditions (*3-5*). Infection by bacterial and viral pathogens has been shown to activate the IRE1α branch of the UPR in immune and epithelial cells, which modulates host responses and disease progression (*6*). Our group further uncovered that IREα signaling drives the inducible expression of *Ptgs2* (Cox-2) and *Ptges* (m-PGES-1) in myeloid cells, promoting prostaglandin biosynthesis and behavioral pain responses in mice (*7*).

During infection, the interplay between host and pathogen factors determines the disease outcome (*8*). Inflammation plays a central role in this relationship and can therefore dictate host survival by affecting the balance between protection and uncontrolled tissue damage (*9*). IRE1α activation has been shown to promote inflammation in various pathophysiological conditions by driving the production of cytokines such as IL-1β, IL-6, IL-23, and TNFα (*10-14*). However, the role of immune-intrinsic IRE1α signaling in the setting of fungal infections is unknown.

Dectin-1 is a C-type lectin receptor that recognizes pathogen-associated molecular patterns (PAMPs) such as β-glucans in fungal cell wall (*15, 16*). Upon binding, the cytoplasmic immunoreceptor tyrosine-based activation motif (ITAM) of Dectin-1 is phosphorylated by the Src family of kinases, thereby providing a docking site for the Spleen Tyrosine Kinase (Syk). Syk subsequently activates protein kinase C-delta (PKCδ) to mediate phosphorylation of the adaptor protein CARD9. This enables CARD9 to complex with BCL10 and MALT1 to activate κβ−dependent transactivation (*16-18*). Syk also relays signals to Vav-PLCγ2, thus activating the NADPH oxidase (NOX) to produce abundant reactive oxygen species (ROS) (*19-21*) that mediate fungal killing (*22, 23*). Recognition of β-glucans by the Dectin1-Syk pathway therefore induces key effector functions in myeloid cells, including phagocytosis, biosynthesis of lipid mediators, ROS production, and expression of cytokines and chemokines such as IL-6, TNFα, IL-1β, CXCL1 and CXCL2, among many others (*24*). Here, we identify a new mechanism whereby fungal β-glucans trigger the IRE1α arm of the UPR in primary neutrophils and monocytes via Dectin1-Src-Syk-NOX activation, ROS production, and the generation of lipid peroxidation byproducts.

*Candida albicans*, normally a commensal fungus, is a human opportunistic pathogen that causes local mucosal infections as well as life-threatening bloodstream systemic infections (*25*). Intravenous *C. albicans* challenge in mice has been extensively used to model clinical disseminated infection (*26*). In this system, *C. albicans* is detected in the brain and kidney after entering the bloodstream, which similarly occurs in humans (*27, 28*). The high blood flow received by the kidney, which reaches 25% of cardiac output, makes this organ a distinctive niche for fungal proliferation, leading to massive recruitment of innate immune cells that aim at controlling the infection at this location (*26, 29, 30*). Nonetheless, unrestrained recruitment of myeloid cells to the infected kidney and the excessive production of inflammatory mediators that accompany this process often cause immune-driven pathology, leading to kidney tissue damage, sepsis, and death (*29-31*). Accordingly, deletion of the chemokine receptor CCR1 or the type-I interferon receptor (IFNAR-1) can reduce neutrophil-driven kidney immunopathology without altering fungal burden in this organ, thus increasing the survival of mice with invasive candidiasis (*32, 33*)

Understanding the immunopathogenic mechanisms of disseminated *C. albicans* infection has been an area of active investigation, but whether immune-intrinsic stress responses are involved in this deadly process has not been established. Here, we uncover that activation of the ER stress sensor IRE1α in myeloid cells promotes the fatal progression of systemic candidiasis in mice. Genetic or pharmacological abrogation of IRE1α reduced kidney immunopathology and extended host survival, suggesting that targeting this pathway might represent a plausible treatment approach for this disease.

## RESULTS

### Systemic *C. albicans* infection triggers IRE1α in kidney-infiltrating immune cells

Myeloid cell activation and accumulation dictate the balance between protective and pathological immune responses during systemic candidiasis (*29, 33-35*), but the involvement of the UPR in controlling this critical balance remains elusive. We evaluated whether systemic infection with *C. albicans* elicited IRE1α activation and the UPR in immune cells *in vivo*. To this end, we used ER stress-activated indicator (ERAI) mice that express the Venus fluorescent protein in cells undergoing IRE1α activation (*36*). C57BL/6J wild type and ERAI mice were intravenously (i.v.) challenged with 10^5^ *C. albicans* SC5314 yeast cells, and their kidneys, blood, spleen, and bone marrow were analyzed at multiple time points thereafter. Consistent with prior reports (*33, 34*), systemic *C. albicans* infection provoked drastic changes in the kidney immune contexture, wherein the proportion of neutrophils and monocytes increased significantly 24- and 72-hours post-infection (**Supplementary Fig. 1, A-C**). These alterations were accompanied by reduced proportions of macrophages, T cells, and B cells in the kidney after infection, while DCs and NK cells remained unchanged (**Supplementary Fig. 1, A-C**). No differences in immune cell infiltration were observed when C57BL/6J wild type or ERAI mice were used as infection hosts (Supplementary Fig. 1, B and C). Notably, neutrophils in the kidney of *C. albicans*-infected mice demonstrated rapid and progressive IRE1α activation, as evidenced by a time-dependent increase in their Venus reporter signal, compared with kidney-resident neutrophils from naïve mice (**Fig. 1, A** and **B**). Blood and spleen neutrophils showed modest IRE1α activation 72 h post-infection, while negligible changes were observed in this population in the bone marrow (**Supplementary Fig. 1, D-F**). Kidney-resident monocytes and DCs in naïve mice demonstrated strong constitutive IRE1α activation, which increased 24 h post-*C. albicans* infection and remained steady thereafter (**Fig. 1, A** and **B)**. Monocytes and DCs at other locations showed minimal changes in IRE1α activation after infection, compared with their naïve counterparts (**Supplementary Fig. 1, D-F**). Of note, systemic challenge with *C. albicans* did not alter IRE1α activation in macrophages, NK cells, T cells, or B cells in the kidney, blood, spleen, or bone marrow at the time points analyzed (**Fig. 1, A** and B; Supplementary Fig. 1, D-F). Therefore, enhanced IRE1α activity is predominantly detected in neutrophils, monocytes, and DCs infiltrating the kidney upon *C. albicans* systemic infection. To confirm these observations, we isolated kidney-resident neutrophils at multiple times post-infection and assessed the expression UPR gene markers. High levels of *Xbp1s, ERdj4, and Grp78* were detected in kidney-infiltrating neutrophils 24 hours after i.v. challenge with *C. albicans*, and the relative abundance of these genes drastically increased in a time-dependent manner (**Fig. 1C**). In these experiments, neutrophils isolated from blood and bone marrow of naïve mice were used as comparative controls since we were unable to isolate enough neutrophils from the kidney of naïve mice. Importantly, despite marked IRE1α and UPR activation in kidney-infiltrating neutrophils from infected mice (**Fig. 1C**), regulated IRE1α-dependent decay (RIDD) was not observed in this setting (**Fig. 1D**). Taken together, these data reveal that systemic candidiasis induces pronounced IRE1α activation in multiple myeloid cell subsets that infiltrate the infected kidney.

**Figure 1.**
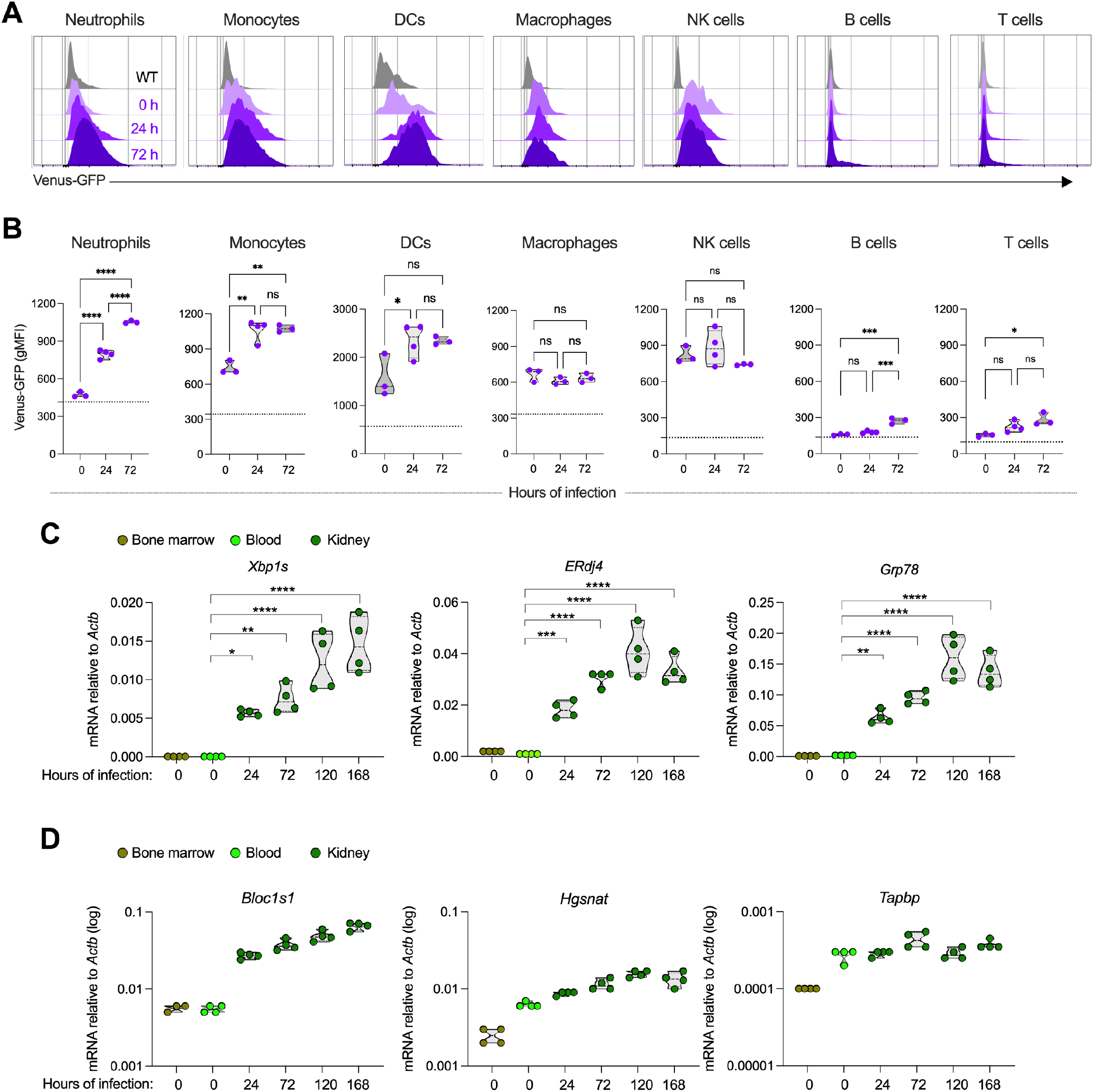
Systemic *C. albicans* infection enhances IRE1α activation in kidney-resident myeloid cells. (**A** and **B)** ERAI or wild type (WT) C57BL/6J mice (*n* = 3-4 per time point) were left untouched or injected i.v. with 10^5^ *C. albicans* cells. Venus reporter expression was assessed by flow cytometry in diverse immune cells infiltrating the kidney at the indicated time points of infection. (**A**) Representative histograms depicting Venus levels in immune cells over time. Gray histograms indicate WT mice that do not express the reporter. Purple histograms denote ERAI mice. (**B**) Geometric mean fluorescence intensity (gMFI) of Venus expression in the indicated immune populations and times. Dashed lines represent intrinsic autofluorescence in WT mice. (**C-D**) WT C57BL/6J mice (*n* = 4 per time point) were injected i.v. with 10^5^ *C. albicans* cells and kidney-infiltrating Ly6G^+^ neutrophils were isolated at days 1, 3, 5 and 7 post-infection. As comparative controls, Ly6G^+^ neutrophils were purified from the blood and bone marrow of naïve mice (*n* = 4). Expression of the indicated transcripts was determined by quantitative RT-PCR and normalized to endogenous *Actb* in each sample. (**C**) UPR-related genes; (**D**) RIDD target genes. Data are shown as mean ± SEM. (**B**) One-way ANOVA (Tukey’s test). (**C**) two-way ANOVA (Dunnett’s test). \**P* < 0.05, \*\**P* < 0.005, \*\*\**P* < 0.0005, \*\*\*\**P* < 0.0001. ns, not significant.

### Dectin-1 engagement induces IRE1α activation via the Src-Syk-NOX pathway

To understand how *C. albicans* instigates ER stress responses in myeloid cells, we used neutrophils as a model system. Neutrophils isolated from the bone marrow of naïve mice were stimulated *ex vivo* with zymosan (a β-glucan-rich Dectin-1 agonist), live *C. albicans*, or bacterial lipopolysaccharide (LPS) that is a TLR4 agonist previously shown to induce IRE1α-XBP1 signaling in various myeloid leukocytes (*7, 11*). Robust IRE1α-dependent splicing of the *Xbp1* mRNA was observed in neutrophils exposed to zymosan or live *C. albicans* (**Fig. 2A**). Real-time quantitative PCR (RT-qPCR) analyses further confirmed that neutrophils exposed to either zymosan or heat-killed *C. albicans* (HKCA) increased the abundance of *Xbp1s* transcripts while markedly upregulating canonical IRE1α/XBP1-target genes such as *Sec61a1* and *ERdj4* (*37*) (**Fig. 2B**), as well as the global UPR marker genes *Grp78, Ddit3*, and *Atf4* (**Fig. 2C**). Hence, zymosan or *C. albicans* stimulation directly triggers ER stress responses in primary neutrophils.

**Figure 2.**
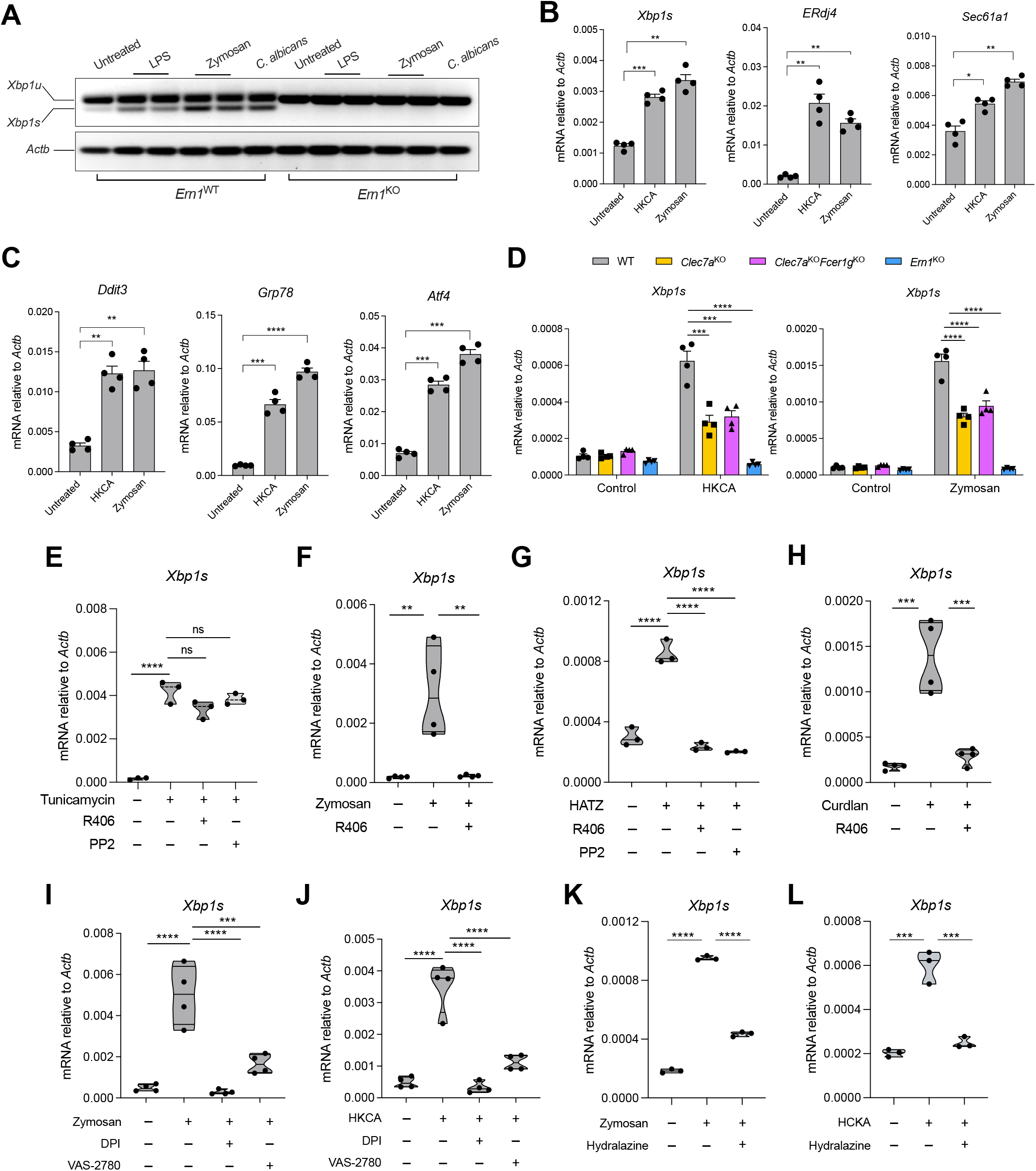
*C. albicans* triggers IRE1α in neutrophils via the Dectin-1-Syk-NOX axis. (**A**) *Ern1*^WT^ or *Ern1*^KO^ neutrophils isolated from the bone marrow were incubated with vehicle control or exposed to LPS (100 ng/ml), zymosan (25 μg/ml), or live *C. albicans* (MOI=10) for 6 hours. *Xbp1* splicing, indicative of IRE1α activation, was evaluated by conventional RT-PCR. *Xbp1u*, unspliced form; *Xbp1s*, spliced form. Data are representative of at least three independent experiments with similar results. (**B** and **C**) Bone marrow-resident neutrophils isolated from WT C57BL/6J mice (*n* = 4) were stimulated with HKCA (MOI=1) or zymosan (25 μg/ml) for 6 hours, and expression of the indicated transcripts was determined by quantitative RT-PCR. (**D**) Bone marrow-resident neutrophils were isolated from mice of the indicated genotypes and then stimulated for 6 hours with HKCA (MOI=10) or zymosan (25 μg/ml). (**E-H**) Dectin-1-mediated activation of IRE1α is driven by Src-Syk. Bone marrow-resident neutrophils from WT C57BL/6J mice (*n* = 3-4) were pretreated for 1 hour with vehicle control or the Syk inhibitor R406 (10 μM), and cells were then stimulated for 6 hours with (**E**) tunicamycin (1 μg/ml), (**F**) zymosan (25 μg/ml), (**G**) hot alkali-treated zymosan (HATZ; 100 μg/ml), or (**H**) curdlan (50 μg/ml). In (**E and G**), neutrophils were also pretreated with the Src family of kinases inhibitor PP2 (10 μM). (**I-L**) Role of NOX-derived ROS and lipid peroxidation byproducts. (**I** and **J**) Bone marrow-resident neutrophils from WT C57BL/6J mice (*n* = 4) were pretreated for 30 min with vehicle control, DPI (10 μM), or VAS-2870 (10 μM) and then stimulated with either (**I**) zymosan (25 μg/ml) or (**J**) HKCA (MOI=10) for 6 hours. (**K** and **L**) Bone marrow-resident neutrophils from WT C57BL/6J mice (*n* = 3) were pretreated for 30 min with vehicle control or hydralazine (100 μg/ml), and then stimulated with either (**K**) zymosan (25 μg/ml) or (**L**) HKCA (MOI=10) for 6 hours. In all cases, *Xbp1s* transcript levels were determined using quantitative RT-PCR, and data was normalized to endogenous *Actb* expression in each sample. Data are shown as mean ± SEM (**B-D**) or violin plots (**E-L**). **B** and **C** two-tailed Student’s *t*-test; (**D-L**) One-way ANOVA (Tukey’s test); \**P* < 0.05, \*\**P* < 0.005, \*\*\**P* < 0.0005, \*\*\*\**P* < 0.0001. ns, not significant. Data are representative of 3-4 independent experiments with similar results.

Dectin-1, encoded by *Clec7a*, is a major C-type lectin receptor (CLR) for β-glucans that enables innate recognition of fungi by host myeloid cells (*38-40*). To determine if activation of IRE1α upon exposure to *C. albicans* yeast cells was mediated by this CLR, we isolated bone marrow-resident neutrophils from wild type or *Clec7a*^KO^ mice and then stimulated them with HKCA or zymosan. In both cases, Dectin-1-deficient neutrophils demonstrated impaired IRE1α activation, as evidenced by a ∼50% reduction in the levels of *Xbp1s* transcripts generated, in comparison with their wild type counterparts (**Fig. 2D**). Similar results were observed when bone marrow-resident monocytes isolated from *Clec7a*^KO^ mice were stimulated with HKCA or zymosan (**Supplementary Fig. 2A**). Additional CLRs such as Dectin-2, Dectin-3, and Mincle can recognize fungal products and induce intracellular signaling via interaction with Fc receptor γ (FcRγ) chain-associated ITAMs (*41*). Thus, to discern whether engagement of these CLRs might contribute to IRE1α activation by *C. albicans*, we tested neutrophils simultaneously lacking both Dectin-1 and FcRγ (*Clec7a*^KO^*Fcer1g*^KO^). A similar decrease in the levels *Xbp1s* was observed in Dectin-1 knockout and Dectin-1/FcRγ double-knockout neutrophils stimulated with either HKCA or zymosan, compared with their wild type counterparts (**Fig. 2D**), suggesting that Dectin-1 is the main CLR mediating IRE1α activation in this setting.

The ITAM of Dectin-1 triggers Syk to induce downstream effector functions such as pathogen killing and cytokine production (*16*). We used the selective Syk inhibitor R406 (*42, 43*) to evaluate whether this kinase mediated Dectin-1-driven IRE1α activation in neutrophils. While R406 did not compromise activation of IRE1α in response to pharmacological ER stress caused by tunicamycin treatment (**Fig. 2E**), this inhibitor completely abrogated *Xbp1* splicing in neutrophils exposed to zymosan (**Fig. 2F**) or hot alkali-treated zymosan (**Fig. 2G**), which abrogates the TLR2-engaging capacity of zymosan while preserving its Dectin-1 agonistic activity (*44*). Similar effects were observed when the Src kinase that operates upstream of Syk was disabled using the pharmacological inhibitor PP2 (*45, 46*) (**Fig. 2G**). In addition, Syk inhibition fully abolished *Xbp1* splicing in neutrophils exposed to curdlan, a bacterial β-glucan that also engages Dectin-1 (**Fig. 2H**). R406 and PP2 also blocked *Xbp1* splicing in primary monocytes exposed to either HKCA or zymosan (**Supplementary Fig. 2B**). Hence, the Src-Syk axis mediates IRE1α activation in neutrophils and monocytes stimulated via Dectin-1.

Upon *C. albicans* or zymosan exposure, Syk activation induces potent ROS generation via NOX (*16, 47, 48*). Importantly, ROS accumulation has been shown to cause ER stress and IRE1α activation by promoting the generation of lipid peroxidation byproducts, such as 4-HNE, that modify ER-resident chaperones and disrupt protein folding in this organelle (*49, 50*). We tested whether NOX-derived ROS could drive IRE1α activation in neutrophils exposed to *C. albicans*. Bone marrow-resident neutrophils stimulated *ex vivo* with either zymosan or HKCA demonstrated copious ROS production, as indicated by a rapid conversion of 2’,7’–dichlorofluorescin diacetate (DCFDA) to its oxidized fluorescent form, 2’,7’–dichlorofluorescein (DCF)(*51*) (**Supplementary Fig. 2, C** and D). Treatment with either diphenyleneiodonium (DPI -an inhibitor of both nitric oxide synthase and NOX) (*52*) or VAS-2870 (a NOX-specific inhibitor) (*53*) suppressed ROS generation (**Supplementary Fig. 2, C**and **D**) and inhibited IRE1α-dependent *Xbp1* splicing (**Fig. 2, I** and **J**) in neutrophils stimulated with zymosan or HCKA. Treatment with Mito-TEMPO or MitoQ that specifically scavenge mitochondrial-derived ROS (*54, 55*) did not affect this process (**Supplementary Fig. 2E**), indicating that NOX-derived ROS are predominant inducers of IRE1α activation in neutrophils stimulated via Dectin-1. Sequestering lipid peroxidation byproducts with the hydrazine derivative hydralazine (*49, 50*) abrogated IRE1α activation in neutrophils sensing zymosan or HCKA (**Fig. 2, K** and **L**). Importantly, treatment with VAS-2870 and hydralazine also prevented IRE1α activation in monocytes sensing zymosan or *C. albicans* (**Supplementary Fig. 2F**). These data uncover that the Dectin-1-Syk-NOX pathway activates ER stress sensor IRE1α in neutrophils and monocytes exposed to *C. albicans* by generating ROS and lipid peroxidation byproducts.

### Selective deletion of IRE1α in leukocytes increases survival in *C. albicans*-infected hosts

We sought to define the role of immune-intrinsic IRE1α during systemic candidiasis. To this end, we used *Ern1*^f/f^*Vav1*^cre^ transgenic mice that selectively delete this ER stress sensor in the hematopoietic compartment (*56, 57*). *Ern1*^f/f^*Vav1*^cre^ mice, or their IRE1α-sufficient (*Ern1*^f/f^) littermate controls, were challenged i.v. with 10^5^ *C. albicans* SC5314 yeast cells and overall host survival was monitored thereafter. Strikingly, loss of IRE1α in leukocytes extended host survival and led to a full recovery in ∼25% of mice with disseminated candidiasis, compared with their IRE1α-sufficient (*Ern1*^f/f^) counterparts (**Fig. 3A**). Notably, deletion of other ER stress sensors, such as PERK or ATF6, did not affect overall host survival after *C. albicans* infection (**Fig. 3, B** and **C**), demonstrating a distinct role for immune-intrinsic IRE1α in the pathogenesis of disseminated candidiasis.

**Figure 3.**
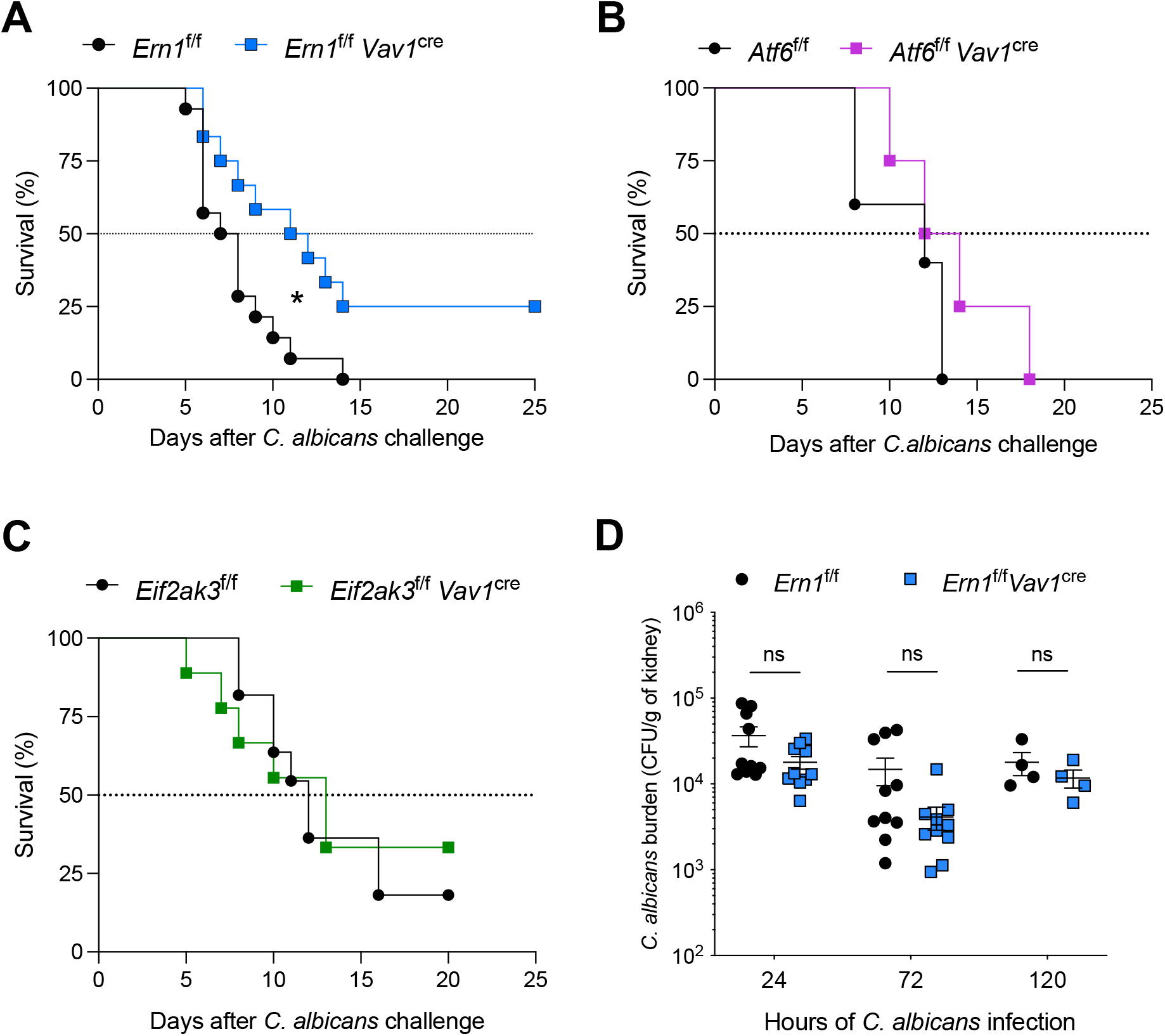
Loss of IRE1α in leukocytes increases survival in mice with systemic candidiasis. (A-C) Mice of indicated genotypes were infected i.v. with 10^5^ *C. albicans*, and host survival was monitored over time. (**A**) *Ern1*^f/f^ (*n* = 14) or *Ern1*^f/f^*Vav1*^cre^ (*n* = 12) mice. (**B**) *Atf6*^f/f^ (*n* = 5) or *Atf6*^f/f^ *Vav1*^cre^ (*n* = 4) mice. (**C**) *Eif2ak3*^f/f^ (*n* = 11) or *Eif2ak3*^f/f^*Vav1*^cre^ (*n* = 9) mice. (**D**) *Ern1*^f/f^ or *Ern1*^f/f^ *Vav1*^cre^ mice (*n* = 4-10 of each genotype per time point) were infected i.v. with 10^5^ *C. albicans*, and fungal burden in the kidney was determined as colony forming units (CFUs) per gram of tissue at the indicated times post-infection. (**A**) Log-rank test; \**P* < 0.05. (**D**) Student’s *t*-test. ns, not significant.

We next evaluated whether ablation of IREα in leukocytes increased host survival by reducing fungal burden in the kidney. Surprisingly, the number of *C. albicans* cells recovered from total kidney homogenates was comparable in mice from both genotypes at multiple days post-infection (**Fig. 3D**), suggesting that loss of IRE1α in leukocytes did not alter *C. albicans killing* in vivo. Indeed, IRE1α-deficient neutrophils, the main effector cells in this setting, showed no alterations in their phagocytic ability (**Supplementary Fig, 3, A** and **B**) or in production of pathogen-killing mediators such as ROS, myeloperoxidase (MPO), or neutrophil extracellular traps (NETs) (**Supplementary Fig. 3, C-H**). Accordingly, the *C. albicans*-killing capacity of IRE1α-deficient neutrophils was comparable to that of their wild type counterparts (**Supplementary Fig. 3I**). Collectively, these data unveil that immune-intrinsic IRE1α signaling supports fatal disease progression in mice systemically infected with *C. albicans*.

### IRE1α promotes kidney tissue damage in mice with systemic *C. albicans* infection

After gaining access to the bloodstream, *C. albicans* can reach the glomeruli and penetrate the renal interstitium through the blood vessels, eliciting an influx of inflammatory myeloid cells capable of causing severe immunopathology and fatal kidney damage (*28, 32, 33*). Thus, we next determined the contribution of IRE1α activation in the immunopathogenesis of invasive candidiasis. *Ern1*^f/f^*Vav1*^cre^ mice, or their IRE1α-sufficient (*Ern1*^f/f^) littermate controls, were challenged i.v. with 10^5^ *C. albicans* SC5314 yeast cells, and their kidneys were resected at different times post-infection for histopathological and immunophenotyping analyses.

Loss of IRE1α in hematopoietic cells did not significantly alter kidney infiltration by monocytes, macrophages, B cells, or NK cells after *C. albicans* challenge (**Supplementary Fig. 4, A-C**). A modest reduction in T cell and DC infiltration was observed in the kidneys of *Ern1*^f/f^*Vav1*^cre^ mice five days after infection, but comparable numbers of these leukocyte subsets were found thereafter in hosts from both genotypes (**Supplementary Fig. 4, E** and **F)**. Of note, the number of total leukocytes and CD45^+^CD11b^+^Ly6G^+^ neutrophils in the kidney were similar in mice from either genotype before infection and also one, three, and five days after *C. albicans* challenge (**Fig. 4, A** and **B**). Yet, we found reduced number of CD45^+^CD11b^+^Ly6G^+^ neutrophils in the kidneys of *Ern1*^f/f^*Vav1*^cre^ mice systemically infected with *C. albicans* for seven days, compared with their IRE1α-sufficient counterparts (**Fig. 4, A** and **B**). Histopathological analyses of kidney sections at this advanced time point confirmed decreased neutrophilic infiltration in *Ern1*^f/f^*Vav1*^cre^ mice, as determined by parallel staining of CD45 and MPO (**Fig. 4, C-F**). Most importantly, hematoxylin and eosin (H&E) staining of the same kidney sections demonstrated that loss of IRE1α in leukocytes significantly diminished the extent of coalescing inflammatory foci while reducing the amount of degenerative and fibrotic lesions, compared with their IRE1α-sufficient counterparts (**Fig. 4, G** and **H**). Hence, immune-intrinsic IRE1α promotes kidney tissue damage in mice with disseminated candidiasis.

**Figure 4.**
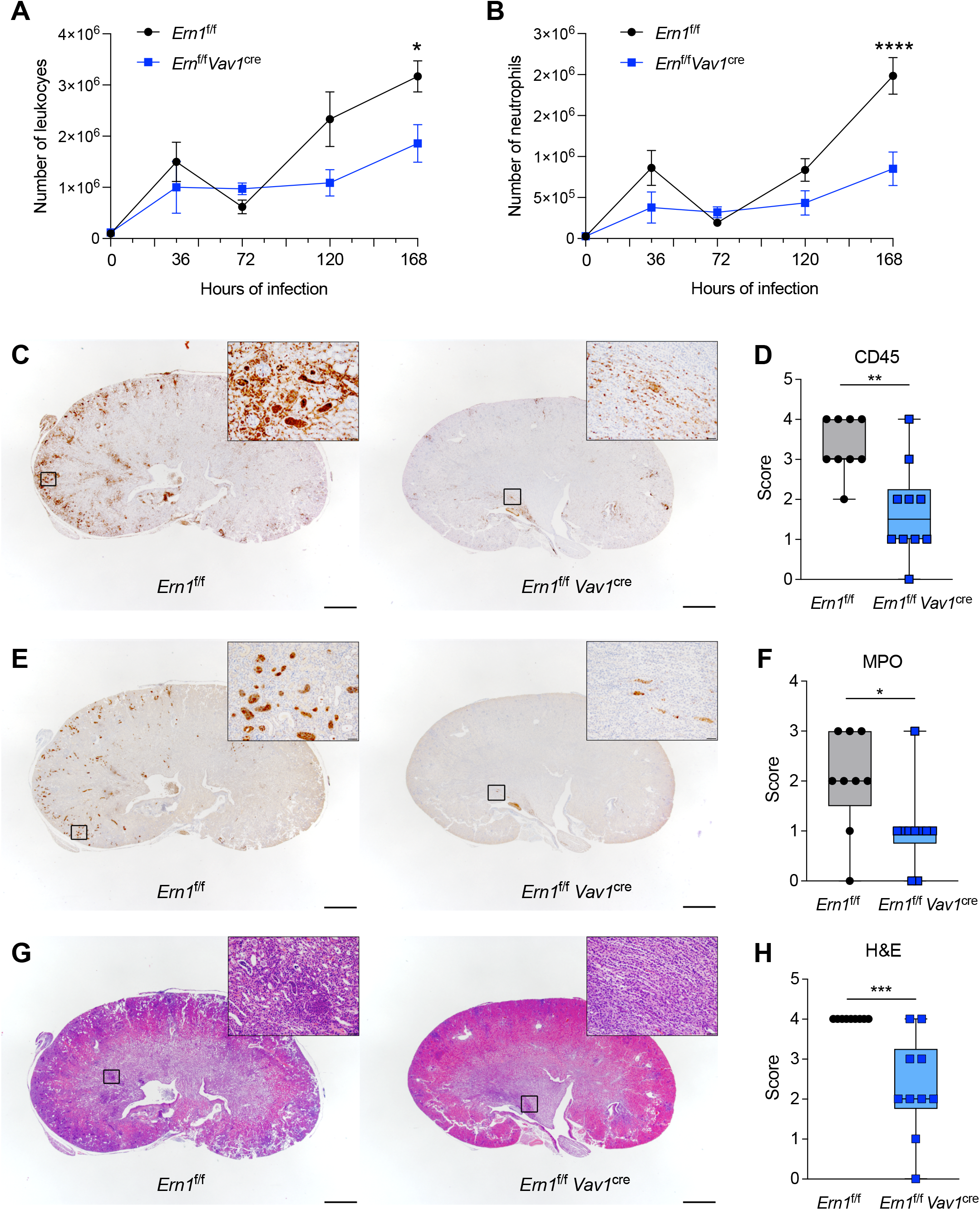
IRE1α deficiency reduces kidney tissue damage in mice with systemic candidiasis. (**A** and **B**) *Ern1*^f/f^ or *Ern1*^f/f^ *Vav1*^cre^ mice (*n* = 3-4 per genotype per time point) were challenged i.v. with 10^5^ *C. albicans* cells and the numbers of CD45^+^ leukocytes (**A**) and CD45^+^CD11b^+^Ly6G^+^Ly6C^low^ neutrophils (**B**) in their kidneys were determined by flow cytometry at the indicated times of infection. (**C-H**) *Ern1*^f/f^ (*n* = 9) or *Ern1*^f/f^ *Vav1*^cre^ (*n* = 10) mice were infected i.v. with 10^5^ *C. albicans* and kidney sections were stained 7 days later for CD45 (**C** and, **D**), MPO (**E** and **F**), and H&E (**G** and **H**). Representative images of kidney section staining (**C, E**, and **G**) and their corresponding pathological scoring (**D, F**, and **H**) are shown. (**A** and **B**) two-way ANOVA (Šídák’s multiple comparisons test). \**P* < 0.05, \*\*\*\**P* < 0.0001. (**D, F**, and **H**) Data are presented as the median plus lower and higher quartiles (boxes) with minimum and maximum values (whiskers). Mann-Whitney test was used for statistical analysis. \**P* < 0.05, \*\**P* < 0.005, \*\*\**P* < 0.0005.

### IRE1α activation drives overt kidney inflammation in hosts with disseminated candidiasis

To further dissect the detrimental effects of IRE1α signaling during systemic *C. albicans* infection, we performed transcriptomic analyses of neutrophils and monocytes sorted from the kidneys of *Ern1*^f/f^ or *Ern1*^f/f^*Vav1*^cre^ mice 36 hours after *C. albicans* i.v. challenge (**Fig. 5A**). Principal component analyses showed a distinct separation of global gene expression profiles based on the status of IRE1α (**Fig. 5B**). We found 190 differentially expressed genes among which 109 were down-regulated and 81 were up-regulated in IRE1α-deficient monocytes and neutrophils, compared with their IRE1α-sufficient counterparts (**Fig. 5, C**and **D**). Of note, downstream pathway analysis using Hallmark gene sets from the Molecular Signature Database (MSigDB) indicated that transcriptional networks related to inflammatory responses, as well as IL6-JAK-STAT3, TNFα, and IL2-STAT5 signaling were significantly downregulated in IRE1α-deficient neutrophils and monocytes infiltrating the kidney, compared with their wild type counterparts (**Fig. 5 E**). Indeed, genes encoding inflammatory mediators such as *Tnf, Il6, Il1a*, and *Ptgs2* were down-regulated in kidney-infiltrating monocytes and neutrophils devoid of IRE1α (**Fig. 5, F**). As expected, loss of IRE1α concomitantly decreased the levels of canonical UPR genes induced by this sensor, such as *Edem1, Sec61a1, Dnajb9* and *Hspa5*, without affecting RIDD (**Fig. 5, F** and **G**).

**Figure 5.**
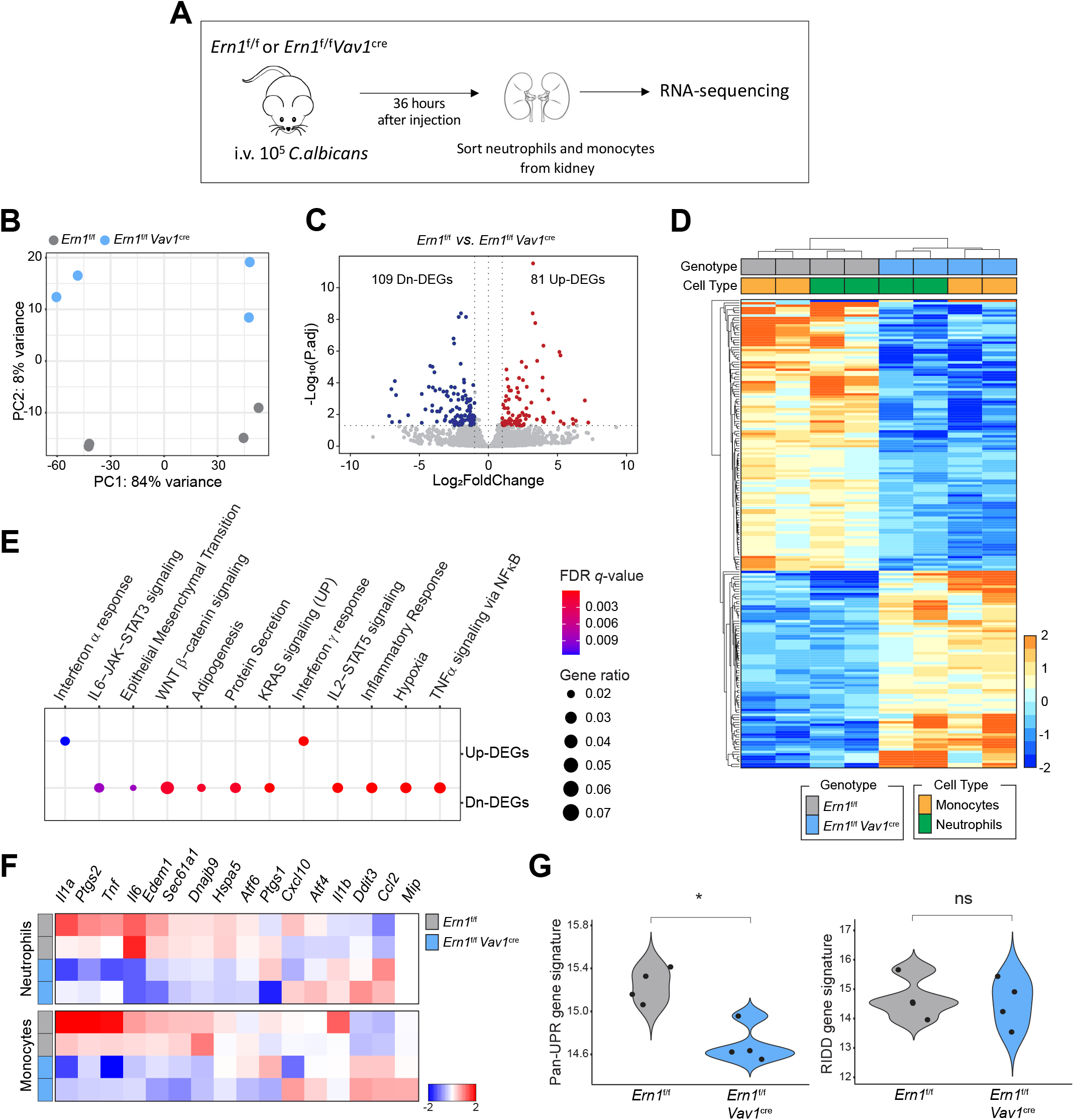
Gene expression profiles controlled by IRE1α in kidney-infiltrating neutrophils and monocytes from mice with systemic candidiasis. (**A-G**) *Ern1*^f/f^ or *Ern1*^f/f^ *Vav1*^cre^ mice (*n* = 4 per genotype) were infected with 10^5^ *C. albicans* cells and 36 hours later, kidney-infiltrating neutrophils (Ly6G^+^ Ly6C^low^) and monocytes (Ly6C^high^) were sorted for RNA-seq analyses. (**A**) Experimental scheme. (**B**) Principal component analysis showing distinct clustering of each genotype. (**C**) Volcano plot describing the abundance of differentially expressed genes (DEGs). Two vertical dashed lines on each side correspond to -1.0 and 1.0 cut-points, which are Log2FoldChange cutoffs used for determining DEGs. The horizontal dashed line corresponds to *P*-adjusted values of 0.05, which was another cutoff used for determining DEGs. (**D**) Heatmap displaying the 81 up-regulated and 109 down-regulated DEGs in *Ern1*^f/f^ *Vav1*^cre^ when compared to *Ern1*^f/f^ cells. (**E**) Pathway enrichment analysis demonstrating down-regulation of various inflammatory pathways. (**F**) Expression profile of select inflammatory and UPR genes in cells of the indicated genotypes. (**G**) Pathway score analyses showing significant downregulation of global UPR gene markers in *Ern1*^f/f^*Vav1*^cre^ neutrophils and monocytes in comparison with their *Ern1*^f/f^ counterparts, whereas no significant difference was observed for RIDD target genes. Wilcoxon test; *\*P* = 0.029; ns, not significant.

High levels of IL-1β, IL-6, and TNFα produced in the kidney at initial stages of systemic *C. albicans* infection have been shown to contribute to the immunopathogenesis the disease (*26, 33, 58*). Importantly, beyond its canonical role in the UPR, IRE1α signaling can enhance the production of diverse inflammatory mediators via multiple mechanisms (*7, 11-13*). We examined whether early IRE1α activation in myeloid cells infiltrating the kidney upon systemic *C. albicans* challenge (**Fig. 1**) promoted the local overexpression of factors contributing to inflammation and renal tissue damage. Thus, we analyzed the levels of multiple cytokines and chemokines in total kidney samples from *Ern1*^f/f^ and *Ern1*^f/f^*Vav1*^cre^ mice systemically infected with *C. albicans* for 3 days, which had similar fungal burden (**Fig. 3D**), as well as comparable immune cell infiltration (**Fig. 4, A** and **B**; **Supplementary Fig. 4, A-F)**. Reduced levels of IL-1β, IL-6, and TNFα were found in supernatants from total kidney homogenates of *Ern1*^f/f^*Vav1*^cre^ mice, compared with their IRE1α-sufficient counterparts at the same time of analysis (**Fig. 6, A-C**). Additional inflammatory mediators such as CCL5, IP-10/CXCL10, IL-1α, MCP-1/CCL2, MIP1α, and PGE_2_ were also significantly decreased in the kidneys of infected *Ern1*^f/f^*Vav1*^cre^ mice at this time point (**Fig. 6, D-I**), while other factors such as IFN-β, IL-10, Eotaxin, IL-2, IL-12 p40, KC, and MCSF remained unaltered (**Supplementary Fig. 5**). Hence, genetic ablation of IRE1α in leukocytes restrains the early overexpression of key pro-inflammatory mediators in the kidney, thereby mitigating the subsequent immune-driven damage to this organ.

**Figure 6.**
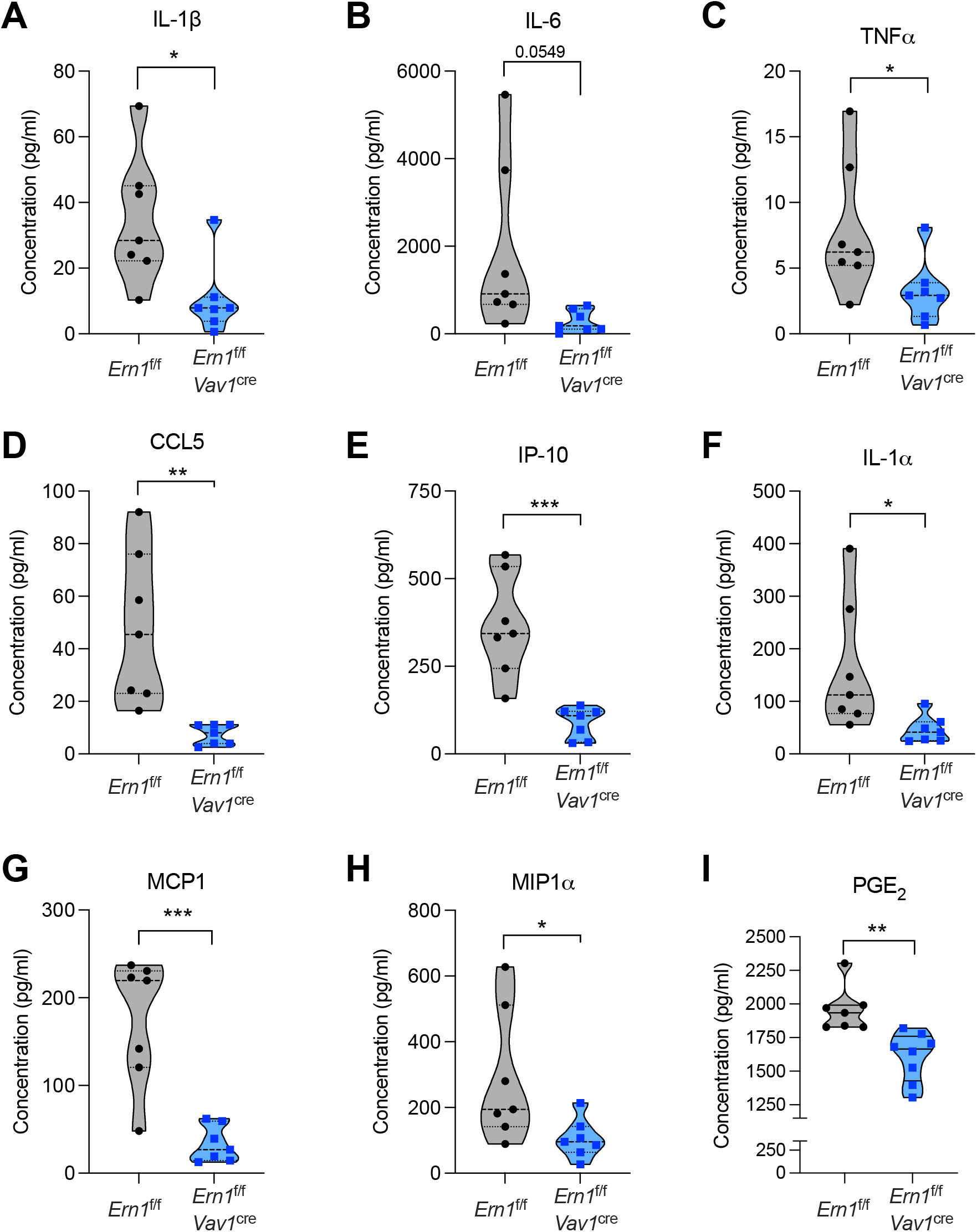
Decreased production of inflammatory factors in the kidneys of *C. albicans*-infected mice lacking IRE1α in leukocytes. (**A-I**) *Ern1*^f/f^ or *Ern1*^f/f^ *Vav1*^cre^ mice (*n* = 7-8 per group) were infected i.v. with 10^5^ *C. albicans* cells and 3 days later, supernatants from total kidney homogenates were analyzed for cytokine and chemokine levels (**A-H**), as well as PGE_2_ expression (**I**). Data are shown using violin plots. Two-tailed Student’s *t*-test was used for statistical analyses. \**P* < 0.05, \*\**P* < 0.005, \*\*\**P* < 0.0005.

### Therapeutic effects of IRE1α inhibition in mice with systemic candidiasis

We next tested whether targeting IRE1α pharmacologically could control kidney inflammation and extend survival in mice systemically infected with *C. albicans*. To this end we used MKC8866, a selective small-molecule inhibitor of IRE1α that has been shown to control the detrimental overactivation of this ER stress sensor in cancer (*59-61*) and inflammatory pain (*7*). Importantly, MKC8866 specifically targets the RNase domain of mammalian IRE1α and does not affect the function of RNase A or RNase L (*61*). *In vitro* treatment with MKC8866 potently inhibited IRE1α activation in primary neutrophils stimulated with either zymosan of HKCA (**Fig. 7, A** and **B**). To test the efficacy of this compound *in vivo*, wild type C57BL/6J mice were challenged i.v. with 10^5^ *C. albicans* SC5314 cells and 24 hours later, mice were treated daily via oral gavage with vehicle control or MKC8866 for 5 consecutive days. Total kidney tissue was analyzed 16 hours after the last dose. MKC8866 administration significantly reduced the levels of IRE1α-generated *Xbp1s* in the kidney of *C. albicans*-infected mice (**Fig. 7C**). Accordingly, the expression of IRE1α/XBP1-dependent target genes in the UPR, such as *ERdj4* and *Sec61a1*, was also significantly decreased in the kidneys from treated mice (**Fig. 7D**), while UPR genes controlled by other ER stress sensors remained unaltered (**Fig. 7E**). Of note, and consistent with our findings using IRE1α-deficient mice (**Fig. 3** and **Fig. 6**), therapeutic MKC8866 administration mitigated the production of IL-1β, TNFα, IL-6, IP-10, CCL5, and PGE_2_ in the kidneys of infected mice (**Fig. 7, F-K**) and extended overall survival, with ∼30% of treated hosts cured of the disease (**Fig. 7L**). Hence, blocking IRE1α activation using a selective small-molecule inhibitor diminishes the immunopathogenic progression of systemic candidiasis in mice.

**Figure 7:**
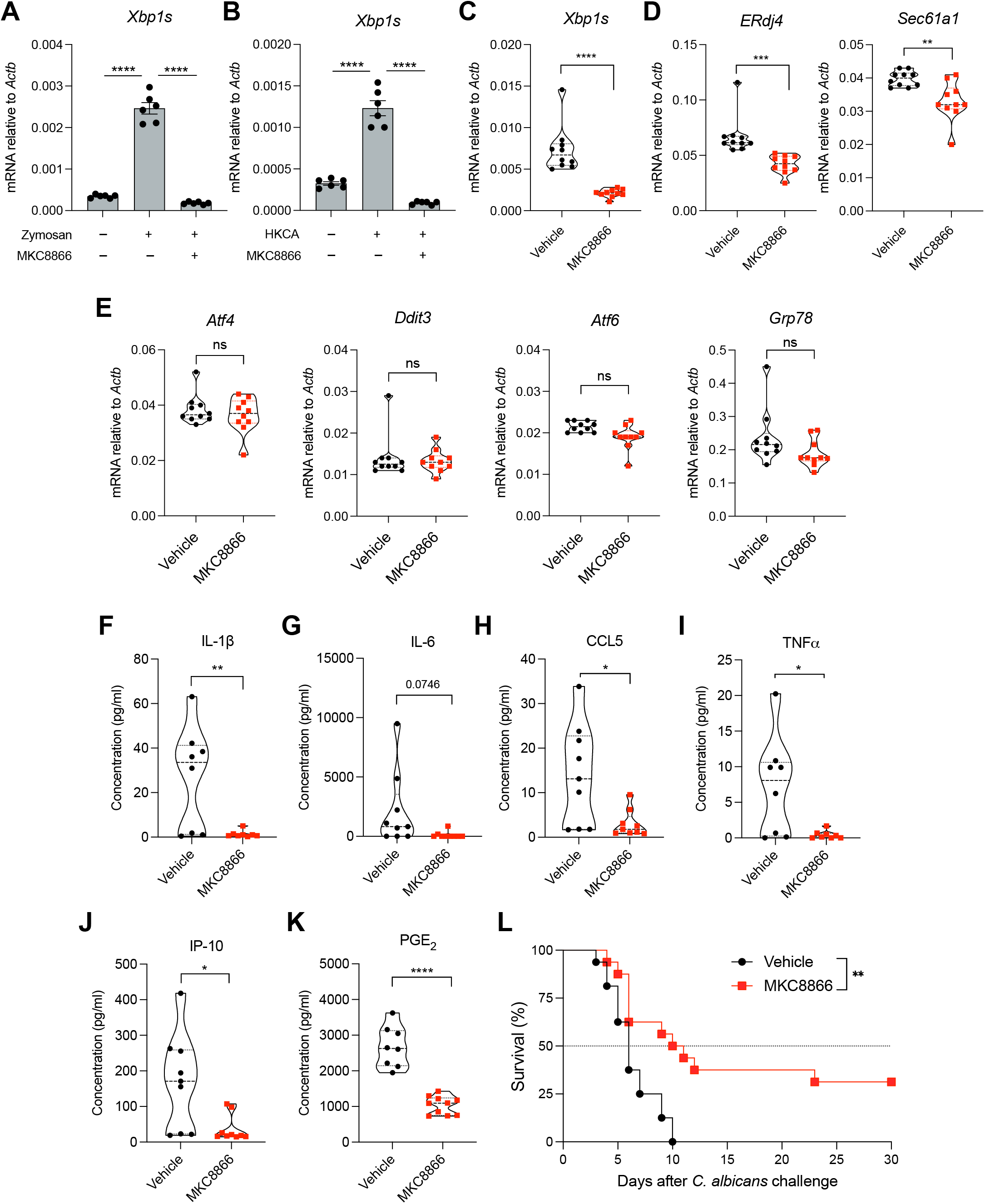
Pharmacological IRE1α inhibition controls kidney inflammation and extends survival in mice with systemic candidiasis. Bone marrow-resident neutrophils from WT C57BL/6J mice were isolated and pretreated for 1 hour with vehicle control or MKC8866 (2.5 μM), followed by stimulation with either **(A)** zymosan (25 μg/ml) or **(B)** HKCA (MOI=10) for 6 hours. *Xbp1s* transcript levels were determined by quantitative RT-PCR and normalized to endogenous *Actb* expression in each sample. (**C-K**) WT C57BL/6J mice (*n* = 8-10 per group) were infected i.v. via tail vein injection with 10^5^ *C. albicans* cells. Twenty-four hours later, mice were treated once daily with vehicle control or MKC8866 (300 mg/kg) via oral gavage until day 5 post-infection, and kidneys were resected 24 hours after the last treatment. (**C-E**) Expression of the indicated transcripts was determined by quantitative RT-PCR and data was normalized to endogenous *Actb* expression in each sample. (**F-K**) Levels of the indicated proinflammatory factors were evaluated in the same kidney samples used in C-E. (**L**) WT C57BL/6J mice (*n* = 16 per group) were infected via tail vein injection with 10^5^ *C. albicans* cells. After 24 hours, mice were treated once daily with vehicle control or MKC8866 (300 mg/kg) via oral gavage for up to 10 days, and host survival was monitored thereafter. (**A** and **B**) Data are shown as mean ± SEM. One-way ANOVA (Tukey’s test); (**C-K**) Two-tailed Student’s *t*-test; (**L**) Log-rank test; \**P* < 0.05, \*\**P* < 0.005, \*\*\**P* < 0.0005, \*\*\*\**P* < 0.0001.

## DISCUSSION

The present study uncovers that inflammatory IRE1α activation promotes lethal immunopathology in the setting of systemic candidiasis. Mechanistically, we determined that sensing of fungal β-glucans by Dectin-1 on myeloid cells triggers robust activation of ER stress sensor IRE1α via ROS generated by the Src-Syk-NOX pathway (**Fig. 8 – proposed model**). The central role of Syk in this process is consistent with its involvement in two additional mechanisms elicited by Dectin-1 that directly impinge on ROS production: the release of arachidonate that activates NOX by acting at multiple steps (*62*) and the generation of the NOX substrate NADPH via the citrate-pyruvate shuttle (*63, 64*). Of note, while treatment with antioxidants fully abrogated IRE1α in neutrophils and monocytes stimulated with zymosan or HCKA, residual activation of this sensor was observed upon genetic loss of Dectin-1. These results suggest that other PRRs, likely TLRs, may contribute to IRE1α activation in this setting via ROS production. Indeed, zymosan β-glucans have been shown to simultaneously activate Dectin-1 and TLR2, while *C. albicans* wall products, such as β-glucans and mannans, can trigger TLR2 and TLR4 concomitantly with Dectin-1 (*65, 66*). Elder and colleagues reported that β-glucans can activate Dectin-1, Syk, NF-kB and p38 signaling, as well as the IRE1α and PERK arms of the UPR in human monocyte-derived DCs, to sustain the expression of thymic stromal lymphopoietin (*67*). Additional studies have shown UPR activation by β-glucans that promotes IL-23 production in macrophages and DCs (*13, 14, 68*). Nonetheless, a detrimental role of leukocyte-intrinsic IRE1α signaling in the setting of systemic candidiasis had not been established.

**Figure 8.**
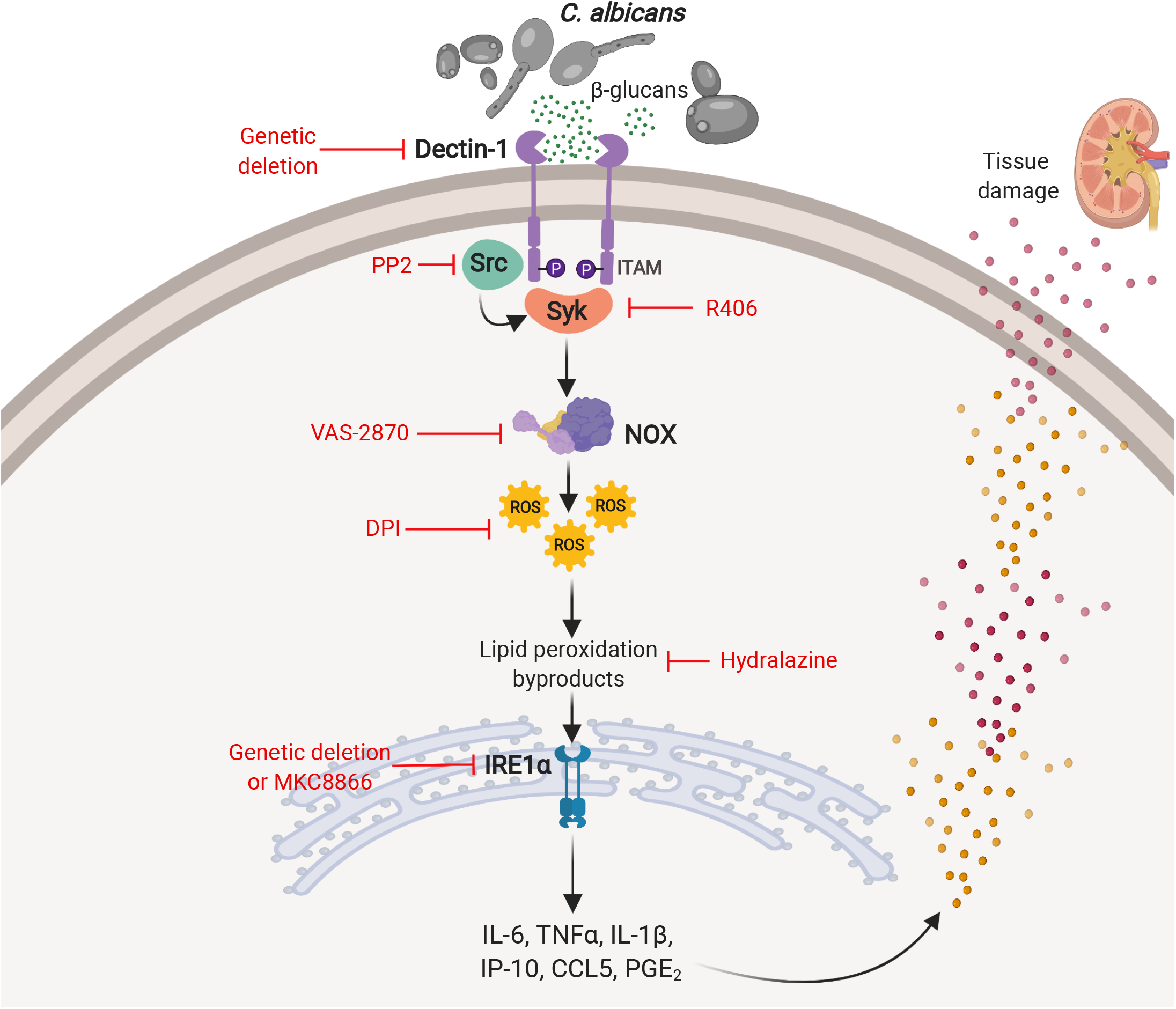
Proposed model: *C. albicans* β-glucans engage Dectin-1 on neutrophils and monocytes, triggering potent Src-Syk-NOX activation. ROS produced by this pathway engender lipid peroxidation byproducts that cause ER stress and fuel IRE1α activation. Robust IRE1α signaling in kidney-resident neutrophils and monocytes of mice with systemic candidiasis mediates the overexpression of multiple inflammatory factors that promote kidney tissue damage and lethal disease progression.

Previous studies identified that host factors, such as the suppressor of TCR signaling 1 and 2 (Sts1/2) (*31*), ubiquitin ligase Casitas C lymphoma-b (CBLB) (*69, 70*), and c-Jun N-terminal Kinase-1 (JNK-1)(*71*), promote lethal immunopathology upon systemic *C. albicans* infection. In addition, global deletion of the chemokine receptor 1 (CCR1) or IFNAR-1 has been shown to reduce immunopathology without affecting fungal burden. In this context, CCR1 selectively regulates the recruitment of neutrophils to the kidney in the late phase of systemic *C. albicans* infection (*33*), while IFNAR-1 dictates the recruitment of Ly6C^hi^ inflammatory monocytes to infected kidneys by driving expression of CCL2 and KC/CXCL1 (*32*). Now, our study unveils that the ER stress sensor IRE1α plays a critical role in the aggressive immunopathogenesis of systemic candidiasis in mice (**Fig. 8**).

We found that loss of IRE1α in the leukocyte compartment controlled the early overproduction of major inflammatory factors, such as IL-1β, IL-1α, IL-6, TNFα, PGE_2_, CCL5, IP-10/CXCL10, and MCP-1/CCL2, in the kidneys of *C. albicans*-infected mice without altering fungal burden. These effects were associated with reduced kidney immunopathology at later stages of infection and increased overall survival, leading to a full recovery in ∼30% of challenged hosts. In addition, our results demonstrated that controlling IRE1α pharmacologically using the small-molecule inhibitor MKC8866 attenuated the overexpression of inflammatory factors in the kidney and significantly extended survival in mice with systemic candidiasis, thus phenocopying the effects of conditional IRE1α deficiency.

In a model of inflammatory placentitis caused by *Brucella abortus*, inhibition of IRE1α blunted IL-6 production and reduced disease severity without affecting bacterial burden (*72*). Similarly, IRE1α was shown to promote inflammation in arthritis and *Francisella* infection models by driving the production of IL-6, TNFα, and IL-1β (*10, 11*). More recent studies have demonstrated that the Sigma-1 receptor inhibits IRE1α activation to control the production of IL-6 and TNFα during bacterial sepsis (*12*), and that IRE1α can enhance the pathogenic activity of immune cells in lupus models (*73*). Here, we propose that IRE1α represents a key target for controlling the detrimental hyperinflammatory condition evoked by systemic infection with *C. albicans*. In patients, invasive candidiasis causes a high mortality rate despite the availability of anti-fungal drugs (*74, 75*). Controlling IRE1α to reduce overt inflammation and tissue damage, while preserving the anti-fungal activity of immune cells, might be useful to manage systemic candidiasis more effectively in the clinic. The role of IRE1α signaling in adaptive immune responses to *C. albicans* in the setting of mucosal, dermal, or vaginal infections (*76*) remains elusive and warrants further investigation.

## ACKNOWLEDGEMENTS

We thank the members of the Flow Cytometry Core Facility and the Genomics Resources Core Facility at Weill Cornell Medicine for their excellent assistance with cell sorting and RNA-seq experiments, respectively. We thank Kim McBride and Maria Jiao for their technical contributions with histologic and immunohistochemical slide preparation. We are grateful to M. Greenblatt, A. Dannenberg, and the members of the Weill Cornell and MSKCC Fungal Interest Group for their comments and suggestions. This work was supported by NIH T32 5T32AI134632-02 and F31CA257631 training grants (A.E.); the Cancer Research Institute-Irvington Institute Postdoctoral Fellowship Award (C-S.C. and C.S.); NIH/NCI Cancer Center Support Grant P30 CA008748 (S.F.S.). NIH R01 NS114653 and R21 CA248106 (E.A.R.-S.); Junta de Castilla y León/Fondo Social Europeo Fellowship (J.J.F.). CSIC’s Global Health Platform (PTI Salud Global, M.S.C.); Plan Nacional de Salud y Farmacia Grant PID2020-113751RB-I00 funded by MCIN/AEI/10.13039/501100011033 (M.S.C.). Junta de Castilla y León/Fondo Social Europeo Grant VA175P20 (M.S.C); NIH R01 DK121977 and the Burroughs Wellcome Fund Investigator in the Pathogenesis of Infectious Diseases Award (I.D.I); NIH R01 093808, NIH R01 139632, NIH R21 142639, and the Burroughs Wellcome Fund Investigator in the Pathogenesis of Infectious Diseases (T.M.H.); NIH R01 NS114653, NIH R21 CA248106, U.S. Department of Defense W81XWH-16-1-0438, OC200166 and OC200224, the Mark Foundation for Cancer Research ASPIRE Award, and The Pershing Square Sohn Foundation (J.R.C-R.)

## METHODS

### Transgenic mice

C57BL/6J, *Vav1*^cre^, *Fcer1g*^-/-^, *Atf6*^f/f^, and *Eif2ak3*^f/f^ mice were obtained from The Jackson Laboratory. ERAI (ER stress-activated indicator) and *Ern1*^f/f^ mice have been previously described by our group (*7, 36, 77*). We generated conditional knockout mice lacking ATF6, PERK, or IRE1α in leukocytes by crossing *Atf6* ^f/f^, *Eif2ak3*^f/f^, or *Ern1* ^f/f^ mice, respectively, with the *Vav1*^cre^ strain that allows selective gene deletion in hematopoietic cells (*7, 56*). *Clec7a*^*-/-*^ mice were generously provided by Dr. Y. Iwakura (*78*). *Clec7a*^*-/-*^*Fcer1g*^*-/-*^ double knockout mice were generated by crossing *Fcer1g*^*-/-*^ mice with *Clec7a*^*-/-*^ mice. All strains used in this study were in the C57BL/6 genetic background. Male and female mice were used at 8-12 weeks of age for all experiments. Mice were maintained in ventilated cages under specific pathogen-free conditions at the animal facilities of Memorial Sloan-Kettering Cancer Center and Weill Cornell Medical College. All *in vivo* experimentation in mice complied with the Weill Cornell Institutional Animal Care and Use Committee (IACUC) under an approved protocol.

### Isolation of neutrophils and monocytes from mouse bone marrow

Untouched neutrophils or monocytes were isolated from the bone marrow of wild type or transgenic mice by negative selection using the Neutrophil Isolation Kit (Miltenyi, catalog #130-097-658) or the Monocyte isolation kit (Miltenyi, catalog #130-100-629), respectively. Briefly, the bone marrow was flushed in complete RPMI media (RPMI + L-glutamine + 10% FBS + HEPES + Sodium Pyruvate + non-essential amino acids + β mercaptoethanol + Pen/strep). Erythrocytes were depleted using ACK lysis buffer and cells were then re-suspended in MACS buffer (0.5% BSA, 1X PBS, 2 mM EDTA). Primary cell isolation was carried out following Miltenyi’s protocols. In all cases, purity was confirmed to be greater than 90% by FACS analysis.

### RNA isolation, quantitative RT-PCR, and *Xbp1* splicing assays

Total RNA was isolated using the RNeasy Mini kit or QIAzol lysis reagent (Qiagen). 0.1-1 μg of total RNA was reverse transcribed to generate cDNA using the qScript cDNA synthesis kit (Quantabio). Quantitative RT-PCR (RT-qPCR) was performed using the PerfeCTa SYBR green fastmix (Quantabio) on a QuantStudio 6 Flex real-time PCR instrument (Applied Biosystems). Normalized gene expression was calculated by comparative threshold cycle method using *Actb* as the endogenous housekeeping gene control. *Xbp1* splicing assays were performed as previously described (*7, 79*). PCR products were separated by electrophoresis through a 2.5% agarose gel and visualized by GelRed staining. All primers used in this study are described in Supplementary Table 1.

### *C. albicans* and β-glucans

We used the *C. albicans* clinical isolate SC5314 (*80*), which was grown in YPD broth (Amresco) for 16 hours at 30°C. Heat-killed *C. albicans* (HKCA) was prepared by heating yeast cultures at 70°C for 30 min. Hot alkali-treated zymosan (HATZ) and curdlan were purchased from Invivogen (Cat # tlrl-zyd and tlrl-curd, respectively). Zymosan was obtained from Sigma-Aldrich (Cat # Z4250-1G). For *in vitro* treatments, neutrophils were stimulated with the agonists described above or co-cultured with live *C. albicans* or HKCA in 96-well plates at 37°C for the indicated time points and MOIs.

### ROS analysis and targeting

Intracellular ROS measurements were performed by labeling with 10 μM 2,7-dichlorofluorescein diacetate (DCFDA) (Thermo-Fischer Scientific, cat# C6827). To define the source of ROS production, 250,000 bone marrow-resident neutrophils were pre-treated for 30 min at 37°C with 10 μM VAS-2870 (Enzo biosciences, cat# BML-E1395-001) or 10 μM DPI (Sigma Aldrich cat# D2926-10MG). Cells were then loaded with DCFDA for 30 min at 37°C and subsequently treated with HKCA, zymosan, or PMA at the indicated MOI or concentrations.

To determine the role of mitochondrial ROS and lipid peroxidation byproducts, bone marrow neutrophils were pre-treated with MitoTEMPO (10 μM, Sigma Aldrich cat# SML0737-5MG), Mitoquinol “MitoQ” (2 μM, Cayman Chemicals # 89950-1MG), or hydralazine (100 μg/ml, Sigma Aldrich # H1753-5G) for 30 min and then stimulated with HKCA or zymosan for the indicated times. After treatment, cells were processed for RNA extraction or flow cytometry analysis.

### Assessment of neutrophil effector functions

Phagocytosis assays were performed by exposing primary neutrophils to FITC-labeled zymosan (Thermo-Fischer Scientific, cat# Z2841) at a 1:1 ratio for 30 minutes at 37°C, followed by a double wash with FACS buffer. Cells were then analyzed on an LSR II flow cytometer. ROS generation was quantified by using DCFDA, as described above. Briefly, 250,000 bone marrow neutrophils from mice of the indicated genotypes were labeled with 10 μM DCDA for 30 min at 37°C. Cells were then stimulated with either HKCA (MOI=5), zymosan (25 μg/ml) or PMA (50 nM, as a positive control) for 1 h at 37°C. Samples were subsequently analyzed on an LSR II flow cytometer. To determine myeloperoxidase (MPO) production, bone marrow neutrophils were isolated from mice of the indicated genotype and stimulated with either HKCA (MOI=5) or zymosan (25 μg/ml) for 6 h in 96-well plates at 37°C. Supernatants were collected after treatment and stored at -80°C until analyzed. Supernatants (25 μl) were used to measure MPO using the Myeloperoxidase Mouse ELISA Kit (Thermo Fisher Scientific, Catalog # EMMPO). Plates were read at 450 nm using a Varioskan Instrument.

DNA release was measured using Sytox Green cell-impermeable nucleic acid stain (Thermo Fisher, cat. no. S7020) as described before(*81, 82*). Bone marrow neutrophils were labeled with 5 μM Sytox Green and then seeded in black 96-well plates at 100,000 cells/well. Cells were subsequently stimulated with HKCA (MOI=20) or zymosan (25 μg/ml) for 8 h. Treatment with PMA (100 nM) was used as a positive control. The fluorescence generated was measured using a Varioskan instrument (Thermo-Fischer Scientific).

Killing assays were performed as described before (*83*). Briefly, primary neutrophils were mixed 1:1 with *C. albicans* yeast cells under gentle agitation at 37°C. 10-μl aliquots were taken at different time points, lysed by re-suspending in water, and fungal CFUs were then determined by serial dilutions on YPD agar plates. Percent killing was evaluated by calculating the number of surviving *C. albicans* in neutrophil co-cultures divided by total *C. albicans* without neutrophils.

### In vivo infection with C. albicans

*C. albicans* SC5314 was cultured as described above and washed three times with 1X PBS. Yeast cells were then counted on a hemocytometer and suspended at a concentration of 5 × 10^5^ cells/ml in PBS. Wild type or transgenic mice of the indicated genotypes were i.v. challenged with 100,000 *C. albicans* SC5314 cells in 200 μl of PBS via tail vein injection. Mice were humanely euthanized at different times post-challenge for downstream analyses or monitored daily to determine overall survival rates.

### Assessment of fungal burden

Mice were humanely euthanized, and kidneys were aseptically removed at the indicated timepoints post-*C. albicans* infection. Kidneys were weighed and ground in 1X PBS with a pestle in 10-cm plates. Homogenates were collected, serially diluted, and plated on YPD agar plates (2% agar) containing chloramphenicol (34 μg/ml) and gentamycin (34 μg/ml). Colony-forming units (CFUs) were determined 48 hours after incubation at 30°C. Fungal burden was calculated as CFU/gram of kidney tissue.

### Analysis of immune cells in kidney, blood, spleen, and bone marrow

Kidney single-cell suspensions were obtained as previously reported (*33*) with some modifications. Briefly, mice were euthanized before or at different time points after infection and kidneys were excised and collected in petri dishes containing 5 ml RPMI + DNase I (0.05mg/ml, Sigma). The kidney was minced on a cell dissociation mesh (Bellcoglass # SKU: 1985-00100) to obtain a single-cell suspension. The suspension was then passed through a 70-μm cell strainer and washed with 20 ml of cold RPMI containing 10% FBS. This cell suspension was centrifuged at 1,500 rpm for 8 min at 4°C. The supernatant was discarded, and red blood cells (RBCs) were lysed using 5 ml ACK lysis buffer for 3 minutes, followed by addition of 20 ml 1X cold PBS. Cell suspensions were immediately passed through a 40-μm cell strainer and again centrifuged at 1,500 rpm for 8 min at 4°C. Supernatants were discarded, and cell pellets were re-suspended in 8 ml of 40% Percoll. This cell suspension was then slowly and carefully overlaid onto 3 ml of 70% Percoll in a 15-ml Falcon tube. The gradient was centrifuged at 2,000 rpm for 30 minutes at room temperature without brakes. The enriched leukocyte fraction was collected from the interphase and washed twice with FACS buffer (1X PBS, 2% FBS and 2 mM EDTA) at 1,500 rpm for 8 min at 4°C. Spleen single-cell suspensions were obtained by directly grinding the spleen onto a 70-μm cell strainer. Bone marrow cells were obtained by flushing the tibiae and femurs of mice with 1X PBS. Circulating leukocytes were analyzed by collecting blood by cardiac puncture. In all cases, RBCs were depleted by treatment with ACK lysis buffer.

Analysis of leukocyte populations in kidney, spleen, blood, and bone marrow single-cell suspensions was performed by flow cytometry using fluorochrome-conjugated antibodies purchased from BioLegend, unless stated otherwise. Cells were Fc-gamma receptor-blocked using TruStain fcXTM (anti-mouse CD16/32, clone 93, Cat# 101319) and then stained for surface markers at 4°C in the dark for 30 minutes with the following antibodies: anti-CD45 (clone 30-F11(RUO) BD Biosciences, Cat# 562420), anti-CD3 (clone 17A2, Cat# 100216), anti-CD19 (clone ID3, BD Biosciences Cat# 612781), anti-CD11b (clone M1/70, Cat# 101257), anti-F4/80 (clone BM8 Cat# 123110), anti-CD11c (clone N418, BD Biosciences Cat# 744180), anti-I-A/I-E (clone M5/114,Cat# 107620), anti-Ly6c (clone HK1.4 Cat# 128041), anti-Ly6G (clone 1A8, Tonbo Biosciences, Cat# 20-1276-U100) and anti-NK1.1 (clone PK136 Cat# 108748). Cells were then washed with 1X PBS and stained with DAPI for live/dead discrimination. Flow cytometry was performed using a Fortessa-X20 instrument (BD Biosciences) and data were analyzed using FlowJo (TreeStar).

### RNA sequencing and bioinformatic analyses

Kidney-infiltrating neutrophils and monocytes were sorted from *Ern1*^f/f^ or *Ern1*^f/f^*Vav1*^cre^ mice 36 h after i.v. challenge with *C. albicans* SC5314. Neutrophils (CD45^+^Ly6g^high^Ly6c^low^CD11b^+^CD3^-^CD19^-^F4/80^-^ CD11c^-^) cells and monocytes (CD45^+^Ly6c^high^CD11b^+^Ly6g^-^CD3^-^CD19^-^F4/80^-^CD11c^-^) cells were sorted on a FACSAria Instrument (BD Biosciences) and processed for RNA isolation using the RNeasy Plus Micro Kit (Qiagen). All samples passed quality controls examined by Agilent Bioanalyzer 2100, and mRNA libraries were generated and sequenced at the Weill Cornell Genomics Resources Core Facility. Raw sequenced reads were pseudoaligned to the mouse reference genome (UCSC mm10) using Kallisto (*84*). Then, transcript abundance was quantified to attain raw counts. Raw counts obtained from Kallisto were used to identify differentially expressed genes (DEGs) using DESeq2 R package (*85*). Genes with very low expression values (baseMean≤15) were filtered out. Then, genes that satisfy two parameters were determined as DEGs: (1) genes with adjusted *P*-value < 0.05 and (2) |Log2FoldChange| > 1.0. Volcano plots were generated to visually display the most significant genes using statistical significance vs. fold-change. To reveal the molecular function of DEGs, Hallmark gene sets from Molecular signature database were used (*86, 87*). The analyses were carried out on the public server (http://www.gsea-msigdb.org/gsea/) (*88*). Single-sample GSEA (ssGSEA) was used to verify whether a given pathway is coordinately up-or down-regulated in a cohort.

ssGSEA computes an enrichment score for each gene set and the score denotes the activity level of a biological process. The analysis was carried out on the public server (https://www.genepattern.org/modules/docs/ssGSEAProjection/4). A Pan-UPR gene set (*Hyou1, Hspa14, Sec61a1, Sec24d, Hspa13, Sec24c, Ambra1, Surf4, Atg13, Xbp1, P4hb, Fam129a, Pdia4, Spcs3, Surf6, Atf6, Dapk1, Dnajb9, Dnajc3, Mfn2, Pdia6, Ccdc47, Dnajc14, Sil1, Sec16a, Tmx3, Sec23b, Sec31a, Gosr2, Asns, Atf4, Ddit3, Ddit4, Edem1*) and *a* RIDD gene set (*Blosc1s1, Hgsnat, Pdgfna, Tapbp, Galnt2, Ergic3, Lamp, Tpp1, Txndc5, Erp44, Rpn1, Idi1, Slc35f5, Pmp22, Col6a1, Pdgfr*) were used for the analyses. [Gene ratio = the number of genes from input that are included in a given gene set / the total number of genes in a given gene set.] For all RNA-seq analyses, statistical significance was set at two-tailed *P*<0.05 and the analyses were conducted using R software (R Foundation for Statistical Computing, Vienna, Austria).

### Histopathology

Following euthanasia, the kidneys were immediately removed and harvested. For each mouse, half kidney was collected and immediately fixed in 10% neutral-buffered formalin. Then, the samples were transferred in 70% ethanol, routinely processed in alcohol and xylene, sectioned at 4 μm thickness, and stained with hematoxylin and eosin (H&E). Immunohistochemistry for myeloperoxidase (Dako A0398, 1:1000, following heat-induced epitope retrieval [HIER] in a pH 6.0 buffer), and CD45 (BD Pharmigen 550539, 1:250, HIER pH6.0) were performed on a Leica Bond RX automated stainer using the Bond Polymer Refine detection system (Leica Biosystem DS9800). The chromogen was 3,3 diaminobenzidine tetrachloride (DAB), and sections were counterstained with hematoxylin.

Specimens were histologically assessed under light microscopy for the detection of lesions. All sections were scored based on the method reported by Wirnsberger et al. with some modifications (*70*). Specifically, the proportion of renal cortex, medulla, and/or pelvis involved by inflammatory changes (tubulointerstitial nephritis and/or pyelonephritis), along with other tubular or interstitial changes were recorded and scored as not significant (score 0), less than 10% tissue affected (score 1), 10-25% tissue affected (score 2), 26-50% tissue affected (score 3), and greater than 50% tissue affected (score 4).

### Analysis of inflammatory factors in kidney samples

Cytokine and chemokine levels in kidneys of mice with systemic *C. albicans* infections were measured by digesting the excised organ in 1 ml of LiberaseTL and DNAse I solution at 37°C for 20 minutes, followed by straining through a 40-μm filter, and centrifugation at 14,000 rpm for 45 minutes at 4°C. Supernatants were collected and stored at -80°C. Cytokine/chemokine analysis in these samples was performed at EVE Technologies (Calgary, AB) using the Mouse Cytokine Array / Chemokine Array 44-Plex (MD44) immunoassay. PGE_2_ levels in kidney supernatants were measured using the PGE_2_ ELISA kit (Enzo Lifesciences, cat# ADI-900-001). Plates for PGE_2_ were read at 405 nm using a Varioskan instrument (Thermo-Fischer Scientific), as we previously reported (*7*).

### *In vivo* treatment with MKC8866

C57BL/6J mice were challenged i.v. with 100,000 *C. albicans* SC5314 yeast cells in 200 μl of PBS via tail vein injection. After 24 hours, infected mice were treated daily via oral gavage with 300 mg/kg MKC8866 (custom manufactured at WuXi AppTec or purchased from MedChemExpress Cat # HY-104040) for the indicated times. The vehicle used for MKC8866 administration was 1% microcrystalline cellulose in 50% sucrose solution, which was prepared by heating at 60°C and stirring until complete solubilization. MKC8866 was added to this vehicle to obtain a final concentration of 30 mg/ml and the mix was sonicated for 1 h at 42 kHz at room temperature to generate a homogeneous suspension. The volume of MKC8866 was given such that each mouse received a dose of 300 mg/kg of body weight.

### Statistical analyses

All statistical analyses were performed using GraphPad Prism 8. Comparison between two groups were assessed using unpaired or paired (for matched comparisons) two-tailed Student’s *t*-test. Mann-Whitney test was used to analyze histopathological scoring. Multiple comparisons were evaluated by one-way ANOVA, including Tukey’s or Dunnett’s multiple comparison tests. Where applicable, error bars represent the mean ± standard error of the mean (SEM). Host survival after *C. albicans* infection was analyzed using the Log-rank (Mantel-Cox) test. *P*-values < 0.05 were considered statistically significant.

---

**Supplementary Figure 1.**
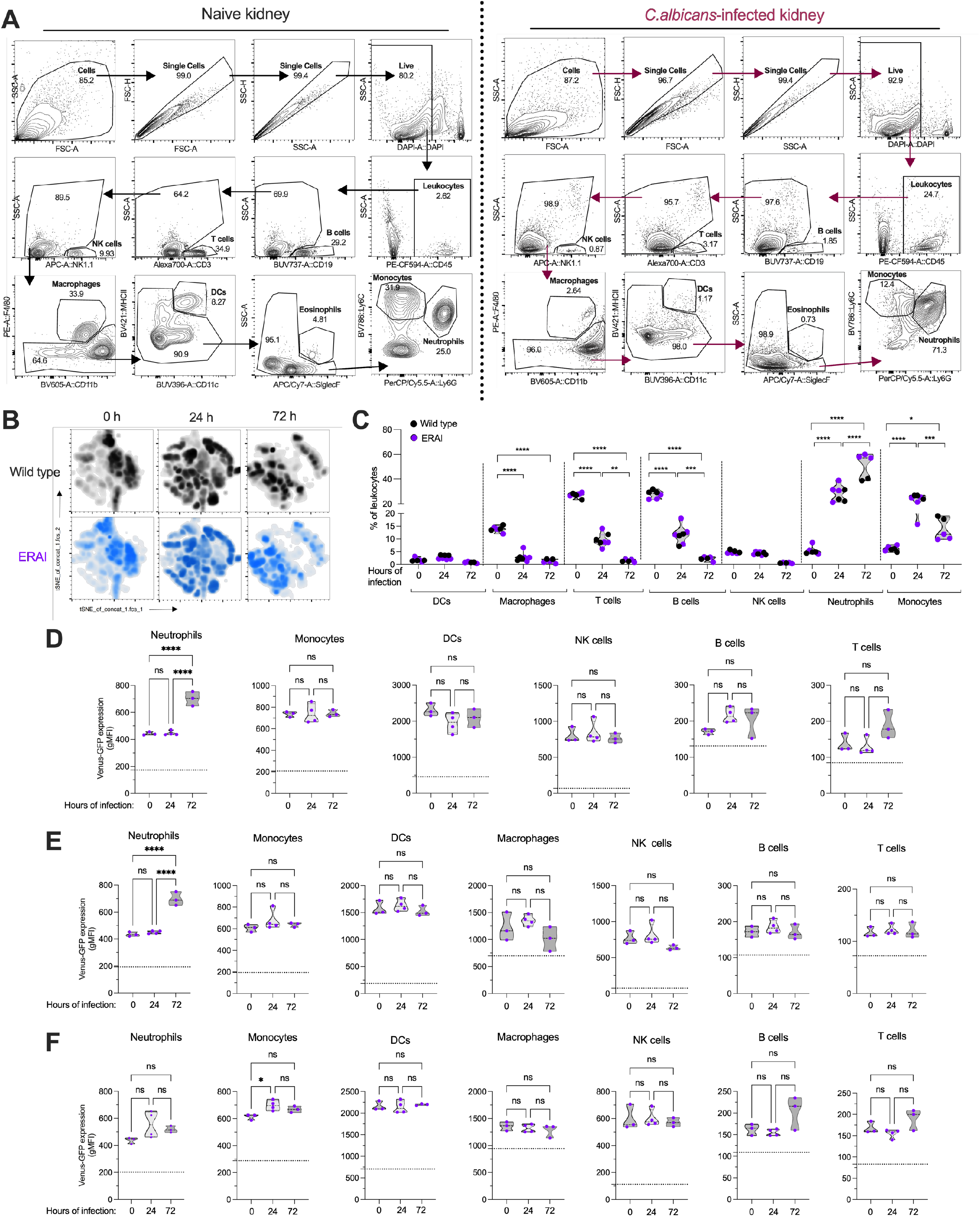
Analysis of WT or ERAI mice with systemic candidiasis. (**A-F**) ERAI or wild type C57BL/6 mice (*n* = 4 per time point) were left untouched or injected i.v. with 10^5^ *C. albicans* cells and their kidneys, blood, spleen, and bone marrow were collected at the indicated time points. Single-cell suspensions were procured from various organs, as described in the methods, and stained with fluorescently-labeled antibodies specific for CD45, CD19, CD3, NK1.1, F4/80, CD11c, MHC-II, SiglecF, CD11b, Ly6C, Ly6G and DAPI. (**A**) Gating strategy used to analyze the kidney immune contexture in naïve or *C. albicans*-infected mice. FACS plots are representative of kidney analysis in an uninfected (naïve) mouse or in a mouse systemically infected with *C. albicans* for 72 h. (**B**) tSNE plots representing time-dependent changes in kidney total CD45^+^ immune cells of ERAI or wild type mice. (**C**) Violin plots showing proportion of the indicated immune cell subsets within total CD45^+^ leukocytes infiltrating the kidney at 0, 24, and 72 h after *C. albicans* infection. (**D-F**) Violin plots for geometric mean fluorescence intensity (gMFI) of Venus reporter expression in the indicated immune cell types in blood (**D**), spleen (**E**), and bone marrow (**F**). Dashed lines represent intrinsic autofluorescence in WT mice. (**C-D**) One-way ANOVA (Tukey’s test). \**P* < 0.05, \*\**P* < 0.005, \*\*\**P* < 0.0005, \*\*\*\**P* < 0.0001, ns, not significant.

**Supplementary Figure 2.**
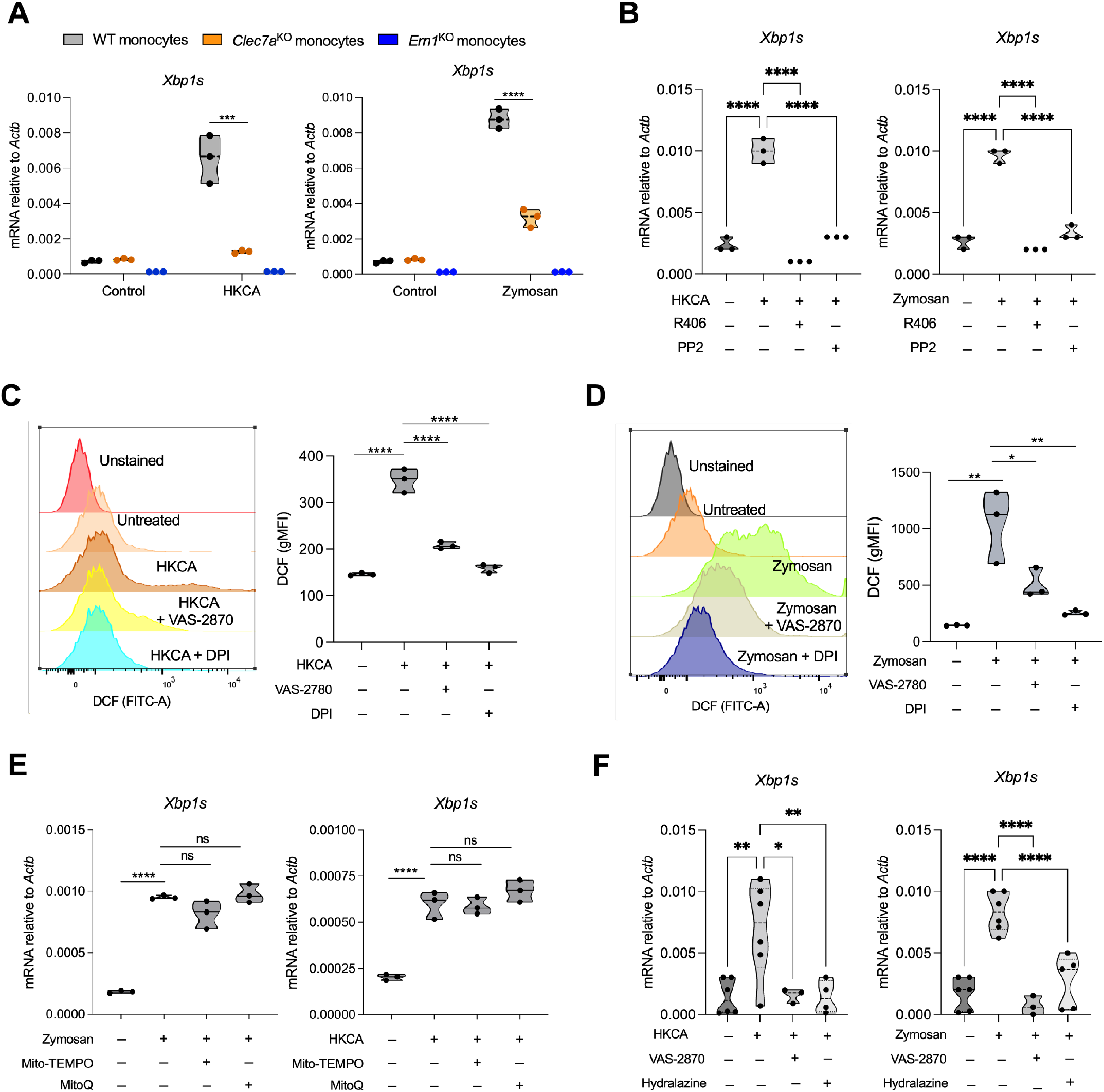
The Dectin-1-Syk-NOX axis also mediates IRE1α activation in monocytes responding to zymosan or *C. albicans*. (**A**) Bone marrow-resident monocytes were isolated from mice of the indicated genotypes and then stimulated for 6 h with HKCA (MOI=10) or zymosan (25 μg/ml). *Xbp1s* transcript levels were measured using quantitative RT-PCR. (**B**) Bone marrow-resident monocytes from WT C57BL/6J mice (*n* = 3) were pretreated for 1 h with vehicle control or the Syk inhibitor R406 (10 μM), and cells were then stimulated for 6 hours with HKCA (MOI=10) or zymosan (25 μg/ml). *Xbp1s* transcript levels were measured using quantitative RT-PCR. **(C** and **D)** WT bone marrow neutrophils (*n* = 3 independent mice) were pretreated for 30 min with vehicle control or ROS inhibitors DPI (10 μM) or VAS-2870 (10 μM) and then stimulated with either **(C)** HKCA (MOI=10) or **(D)** zymosan (25 μg/ml) for 1 hour. ROS production was measured by flow cytometry as described in the methods. (**E**) WT bone marrow neutrophils (*n* = 3 independent mice) were pretreated for 30 min with vehicle control or mitochondrial ROS scavengers, Mito-TEMPO (10 μM) or MitoQ (2 μM), and then stimulated with either zymosan (25 μg/ml) or HKCA (MOI=10) for 6 hours. *Xbp1s* transcript levels were measured using quantitative RT-PCR. (**F**) Bone marrow-resident monocytes isolated from WT mice were pretreated with VAS-2870 or hydralazine and then stimulated for 6 hours with HKCA (MOI=10) or zymosan (25 μg/ml). *Xbp1s* transcript levels were measured using quantitative RT-PCR. Data are shown as violin plots. One-way ANOVA (Tukey’s test) was used for statistical analysis; \**P* < 0.05, \*\**P* < 0.005, **** *P* <0.0001. gMFI, geometric mean fluorescence intensity. ns, not significant.

**Supplementary Figure 3.**
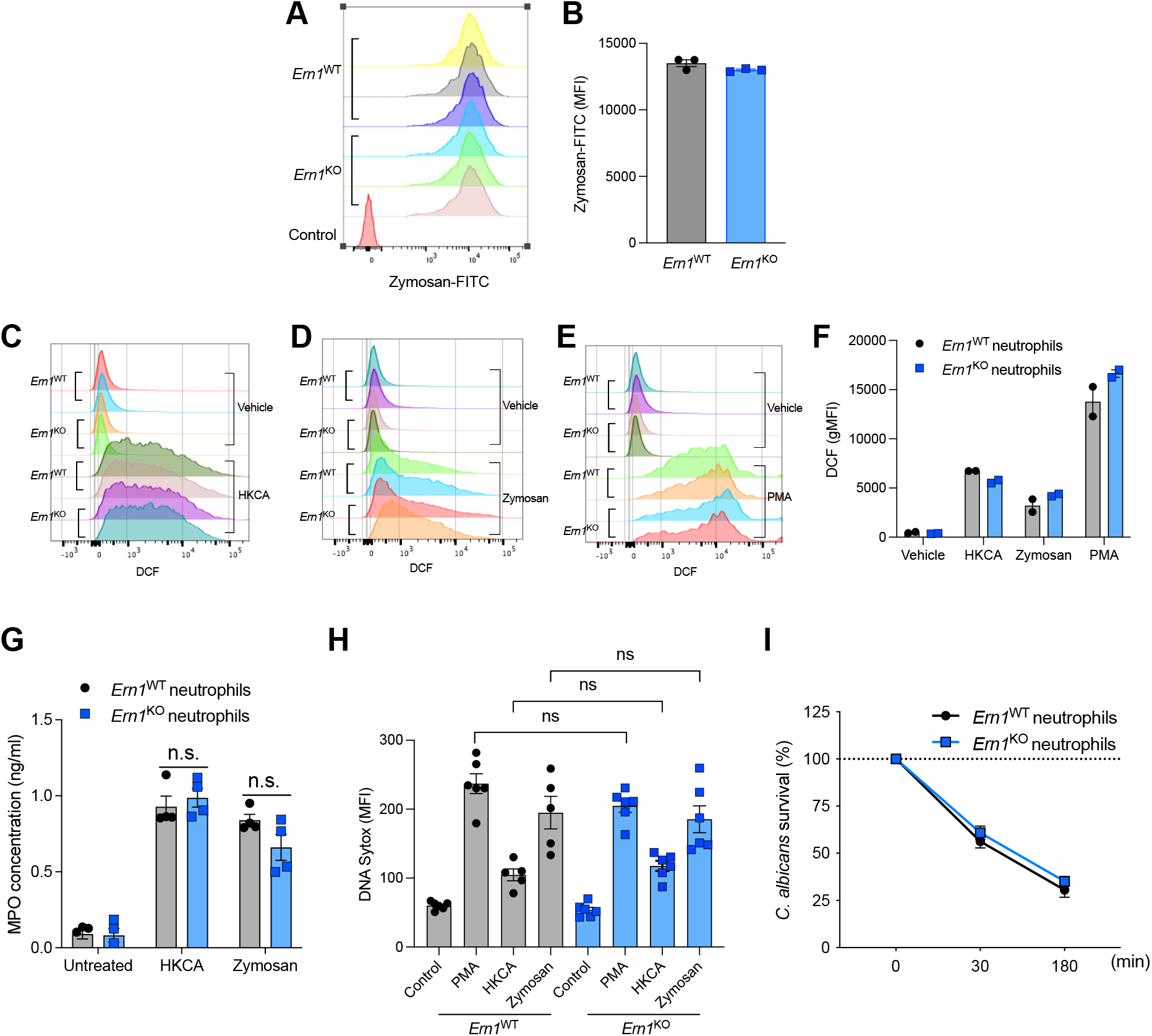
Loss of IRE1α does not alter anti-*C. albicans* effector functions in neutrophils. (**A** and **B**) *Ern1*^WT^ or *Ern1*^KO^ neutrophils were incubated with FITC-labeled zymosan for 30 min and their phagocytic capacity was assessed by FACS. (**C-F**) *Ern1*^WT^ or *Ern1*^KO^ neutrophils were stimulated with (**C**) HKCA (MOI=5), (**D**) zymosan (25 μg/ml) or (**E**) PMA (50 nM) for 1 h. ROS production was then quantified by FACS using the intensity of DCF signal generated (**F**). **(G)** *Ern1*^WT^ or *Ern1*^KO^ neutrophils were stimulated with HKCA (MOI=5) or zymosan (25 μg/ml) for 6 h and myeloperoxidase (MPO) production was measured in supernatants by ELISA. **(H)** *Ern1*^WT^ or *Ern1*^KO^ neutrophils were stimulated with HKCA (MOI=20), zymosan (25 μg/ml) or PMA (100 nM) for 8 hours and DNA release was measured as a marker for NETosis using Sytox green. (**I**) *Ern1*^WT^ or *Ern1*^KO^ neutrophils (*n* = 4) were isolated and co-cultured with the yeast form of *C. albicans* for the indicated time points and CFUs were determined by serial dilutions on YPD agar. Percent survival was determined by normalization to *C. albicans* cultured without neutrophils. Data are shown as mean ± SEM. One-way ANOVA (Tukey’s test) was used for statistical analysis; ns, not significant; MFI, mean fluorescence intensity; gMFI, geometric mean fluorescence intensity.

**Supplementary Figure 4.**
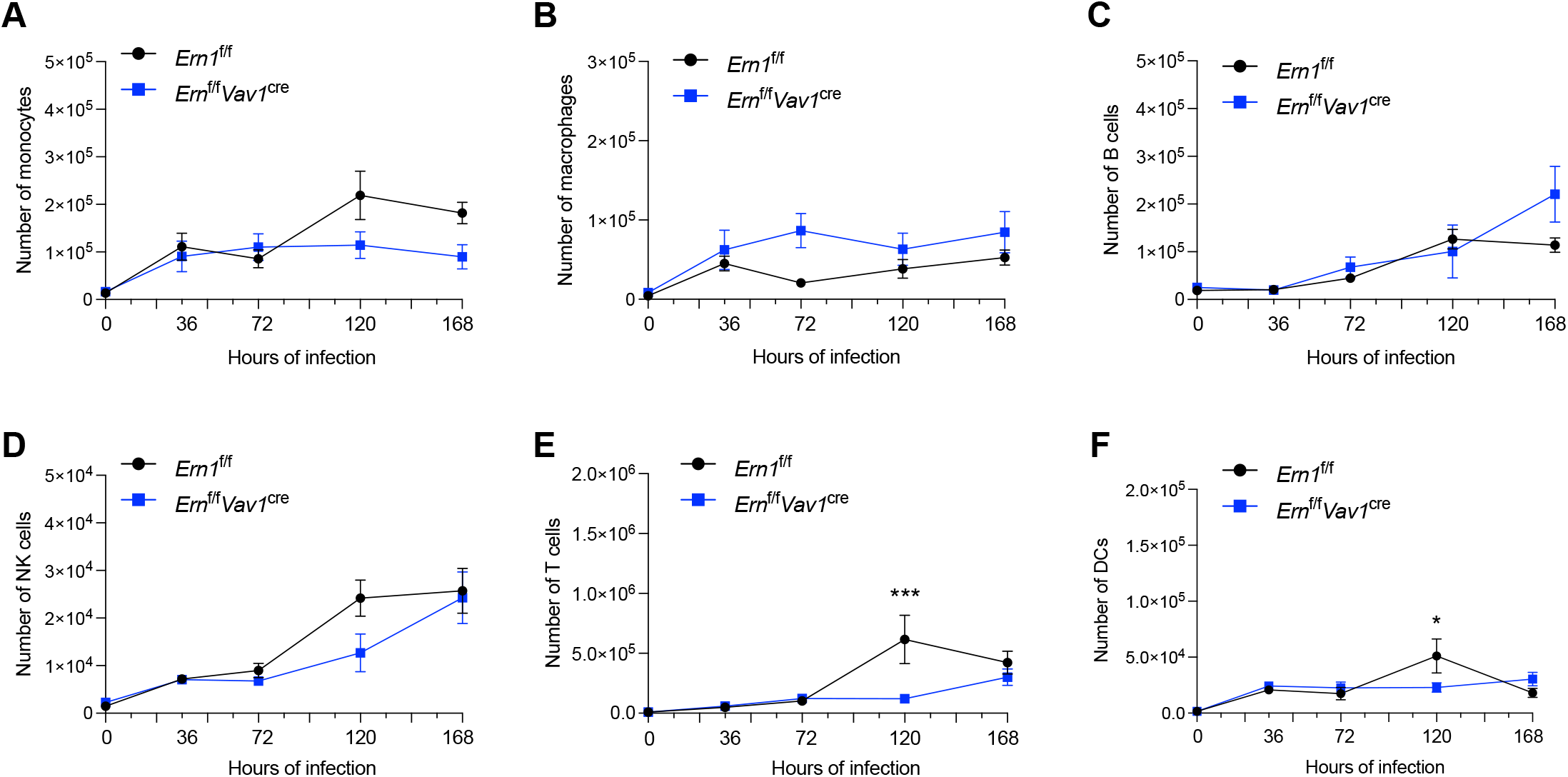
Additional immune cells infiltrating the kidney during systemic *C. albicans-*infection. *Ern1*^f/f^ or *Ern1*^f/f^ *Vav1*^cre^ mice (*n* = 3-4 per genotype per time point) were infected i.v. with 10^5^ *C. albicans* cells and the number of (**A**) monocytes, (**B**) macrophages, (**C**) B cells, (**D**) NK cells, (**E**) T cells, and (**F**) DCs in the kidney were determined by flow cytometry at the indicated times. Data are shown as mean ± SEM. Two-way ANOVA (Šídák’s multiple comparisons test). \**P* < 0.05, \*\*\**P* < 0.0005.

**Supplementary Figure 5.**
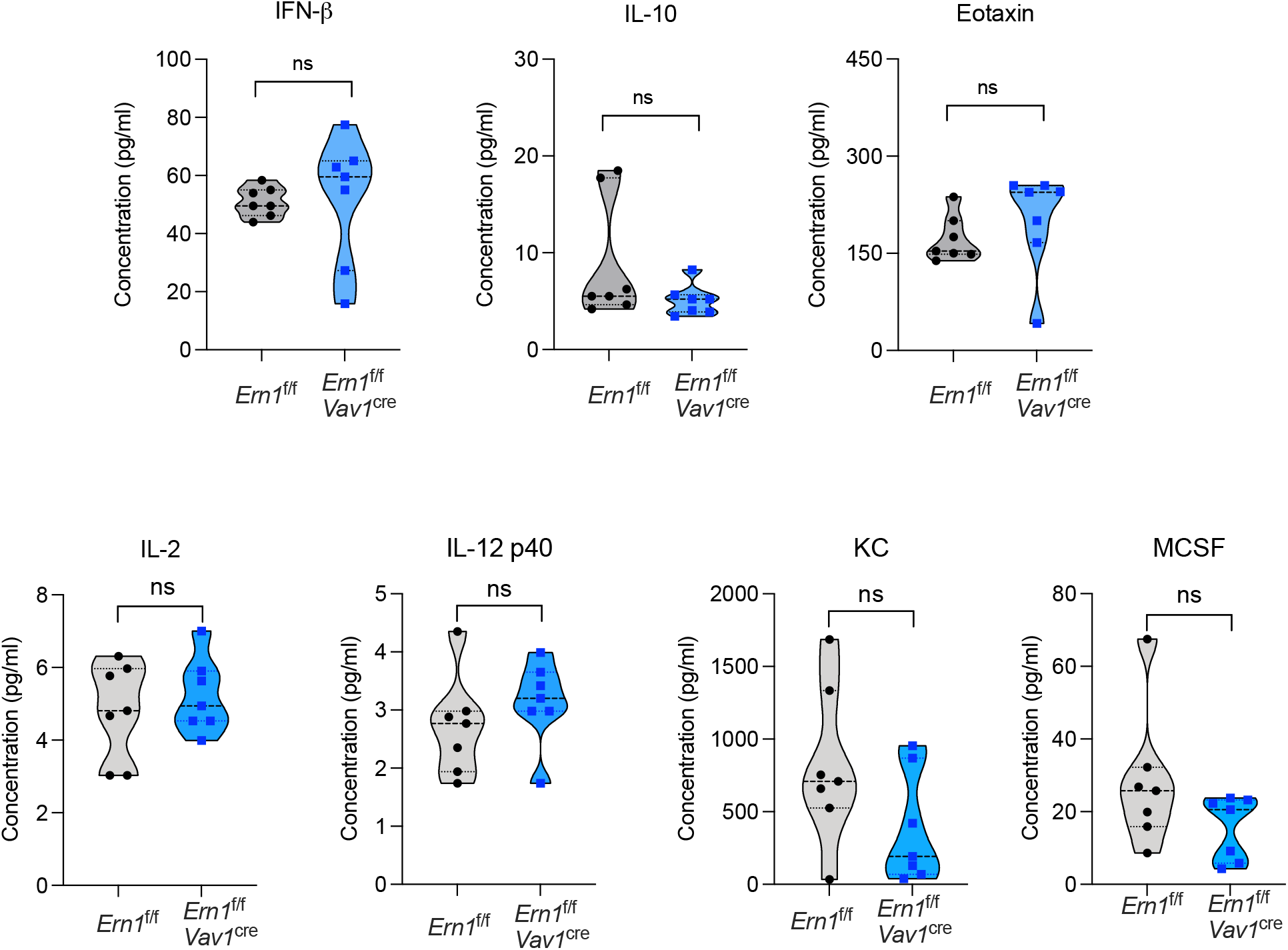
Additional cytokines in kidney homogenates from *C. albicans-*infected mice. (**A-C**) *Ern1*^f/f^ (n=7) or *Ern1*^f/f^ *Vav1*^cre^ (n=7) mice were infected i.v. with 10^5^ *C. albicans* and expression of the indicated factors was determined by ELISA in total kidney homogenates 3 days post-infection. Data are shown as violin plots with dots representing independent mice. Two-tailed Student’s *t*-test was used for statistical analysis. ns, not significant.

**Supplementary Table 1.**
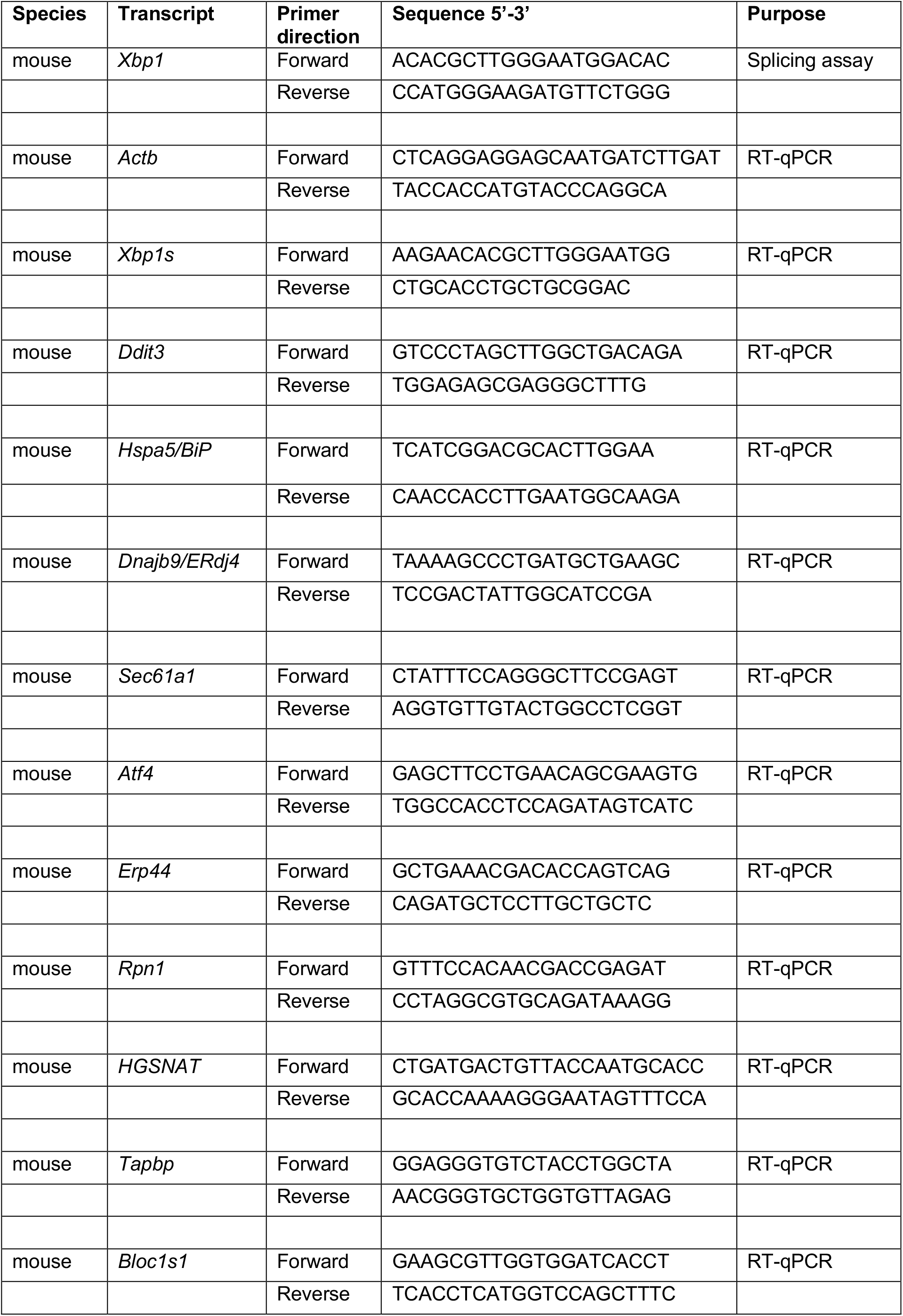
Primers used in this study.

## REFERENCES

1. C. Hetz, K. Zhang, R. J. Kaufman, Mechanisms, regulation and functions of the unfolded protein response. Nat Rev Mol Cell Biol, (2020).

2. J. Celli, R. M. Tsolis, Bacteria, the endoplasmic reticulum and the unfolded protein response: friends or foes? Nat Rev Microbiol 13, 71–82 (2015).

3. S. E. Bettigole, L. H. Glimcher, Endoplasmic reticulum stress in immunity. Annu Rev Immunol 33, 107–138 (2015).

4. X. Chen, J. R. Cubillos-Ruiz, Endoplasmic reticulum stress signals in the tumour and its microenvironment. Nat Rev Cancer 21, 71–88 (2021).

5. C. Hetz, J. M. Axten, J. B. Patterson, Pharmacological targeting of the unfolded protein response for disease intervention. Nature chemical biology 15, 764–775 (2019).

6. J. A. Choi, C. H. Song, Insights Into the Role of Endoplasmic Reticulum Stress in Infectious Diseases. Frontiers in immunology 10, 3147 (2019).

7. S. Chopra et al., IRE1alpha-XBP1 signaling in leukocytes controls prostaglandin biosynthesis and pain. Science 365, (2019).

8. A. Casadevall, L. A. Pirofski, Host-pathogen interactions: basic concepts of microbial commensalism, colonization, infection, and disease. Infect Immun 68, 6511–6518 (2000).

9. M. A. Jabra-Rizk et al., Candida albicans Pathogenesis: Fitting within the Host-Microbe Damage Response Framework. Infect Immun 84, 2724–2739 (2016).

10. Q. Qiu et al., Toll-like receptor-mediated IRE1alpha activation as a therapeutic target for inflammatory arthritis. EMBO J 32, 2477–2490 (2013).

11. F. Martinon, X. Chen, A. H. Lee, L. H. Glimcher, TLR activation of the transcription factor XBP1 regulates innate immune responses in macrophages. Nat Immunol 11, 411–418 (2010).

12. D. A. Rosen et al., Modulation of the sigma-1 receptor-IRE1 pathway is beneficial in preclinical models of inflammation and sepsis. Sci Transl Med 11, (2019).

13. S. Marquez et al., Endoplasmic Reticulum Stress Sensor IRE1alpha Enhances IL-23 Expression by Human Dendritic Cells. Frontiers in immunology 8, 639 (2017).

14. D. A. Mogilenko et al., Metabolic and Innate Immune Cues Merge into a Specific Inflammatory Response via the UPR. Cell 177, 1201–1216 e1219 (2019).

15. M. J. Marakalala, A. M. Kerrigan, G. D. Brown, Dectin-1: a role in antifungal defense and consequences of genetic polymorphisms in humans. Mamm Genome 22, 55–65 (2011).

16. A. Verma, M. Wuthrich, G. Deepe, B. Klein, Adaptive immunity to fungi. Cold Spring Harb Perspect Med 5, a019612 (2014).

17. R. A. Drummond, G. D. Brown, The role of Dectin-1 in the host defence against fungal infections. Curr Opin Microbiol 14, 392–399 (2011).

18. H. Yang, H. He, Y. Dong, CARD9 Syk-dependent and Raf-1 Syk-independent signaling pathways in target recognition of Candida albicans by Dectin-1. Eur J Clin Microbiol Infect Dis 30, 303–305 (2011).

19. D. B. Graham et al., Neutrophil-mediated oxidative burst and host defense are controlled by a Vav-PLCgamma2 signaling axis in mice. J Clin Invest 117, 3445–3452 (2007).

20. B. H. Segal, M. J. Grimm, A. N. Khan, W. Han, T. S. Blackwell, Regulation of innate immunity by NADPH oxidase. Free Radic Biol Med 53, 72–80 (2012).

21. A. Mocsai, M. Zhou, F. Meng, V. L. Tybulewicz, C. A. Lowell, Syk is required for integrin signaling in neutrophils. Immunity 16, 547–558 (2002).

22. S. Saijo, Y. Iwakura, Dectin-1 and Dectin-2 in innate immunity against fungi. Int Immunol 23, 467–472 (2011).

23. G. T. Nguyen, E. R. Green, J. Mecsas, Neutrophils to the ROScue: Mechanisms of NADPH Oxidase Activation and Bacterial Resistance. Front Cell Infect Microbiol 7, 373 (2017).

24. D. M. Reid, N. A. Gow, G. D. Brown, Pattern recognition: recent insights from Dectin-1. Curr Opin Immunol 21, 30–37 (2009).

25. F. L. Mayer, D. Wilson, B. Hube, Candida albicans pathogenicity mechanisms. Virulence 4, 119–128 (2013).

26. D. M. MacCallum, Massive induction of innate immune response to Candida albicans in the kidney in a murine intravenous challenge model. FEMS Yeast Res 9, 1111–1122 (2009).

27. J. C. Parker, Jr., J. J. McCloskey, K. A. Knauer, Pathobiologic features of human candidiasis. A common deep mycosis of the brain, heart and kidney in the altered host. Am J Clin Pathol 65, 991–1000 (1976).

28. C. V. Jawale, P. S. Biswas, Local antifungal immunity in the kidney in disseminated candidiasis. Curr Opin Microbiol 62, 1–7 (2021).

29. M. S. Lionakis, J. K. Lim, C. C. Lee, P. M. Murphy, Organ-specific innate immune responses in a mouse model of invasive candidiasis. J Innate Immun 3, 180–199 (2011).

30. B. Spellberg, A. S. Ibrahim, J. E. Edwards, Jr., S. G. Filler, Mice with disseminated candidiasis die of progressive sepsis. J Infect Dis 192, 336–343 (2005).

31. S. Naseem, D. Frank, J. B. Konopka, N. Carpino, Protection from systemic Candida albicans infection by inactivation of the Sts phosphatases. Infect Immun 83, 637–645 (2015).

32. O. Majer et al., Type I interferons promote fatal immunopathology by regulating inflammatory monocytes and neutrophils during Candida infections. PLoS Pathog 8, e1002811 (2012).

33. M. S. Lionakis et al., Chemokine receptor Ccr1 drives neutrophil-mediated kidney immunopathology and mortality in invasive candidiasis. PLoS Pathog 8, e1002865 (2012).

34. J. V. Desai, M. S. Lionakis, The role of neutrophils in host defense against invasive fungal infections. Curr Clin Microbiol Rep 5, 181–189 (2018).

35. M. G. Netea, L. A. Joosten, J. W. van der Meer, B. J. Kullberg, F. L. van de Veerdonk, Immune defence against Candida fungal infections. Nat Rev Immunol 15, 630–642 (2015).

36. T. Iwawaki, R. Akai, K. Kohno, M. Miura, A transgenic mouse model for monitoring endoplasmic reticulum stress. Nat Med 10, 98–102 (2004).

37. A. H. Lee, N. N. Iwakoshi, L. H. Glimcher, XBP-1 regulates a subset of endoplasmic reticulum resident chaperone genes in the unfolded protein response. Mol Cell Biol 23, 7448–7459 (2003).

38. G. D. Brown, Dectin-1: a signalling non-TLR pattern-recognition receptor. Nat Rev Immunol 6, 33–43 (2006).

39. H. S. Goodridge, A. J. Wolf, D. M. Underhill, Beta-glucan recognition by the innate immune system. Immunol Rev 230, 38–50 (2009).

40. G. D. Brown et al., Dectin-1 mediates the biological effects of beta-glucans. J Exp Med 197, 1119–1124 (2003).

41. J. Tang, G. Lin, W. Y. Langdon, L. Tao, J. Zhang, Regulation of C-Type Lectin Receptor-Mediated Antifungal Immunity. Frontiers in immunology 9, 123 (2018).

42. S. Braselmann et al., R406, an orally available spleen tyrosine kinase inhibitor blocks fc receptor signaling and reduces immune complex-mediated inflammation. J Pharmacol Exp Ther 319, 998–1008 (2006).

43. P. Ruzza, B. Biondi, A. Calderan, Therapeutic prospect of Syk inhibitors. Expert Opin Ther Pat 19, 1361–1376 (2009).

44. B. N. Gantner, R. M. Simmons, S. J. Canavera, S. Akira, D. M. Underhill, Collaborative induction of inflammatory responses by dectin-1 and Toll-like receptor 2. J Exp Med 197, 1107–1117 (2003).

45. K. R. Brandvold, M. E. Steffey, C. C. Fox, M. B. Soellner, Development of a highly selective c-Src kinase inhibitor. ACS Chem Biol 7, 1393–1398 (2012).

46. J. H. Hanke et al., Discovery of a novel, potent, and Src family-selective tyrosine kinase inhibitor. Study of Lck- and FynT-dependent T cell activation. J Biol Chem 271, 695–701 (1996).

47. O. Gross et al., Syk kinase signalling couples to the Nlrp3 inflammasome for anti-fungal host defence. Nature 459, 433–436 (2009).

48. Y. S. Bae et al., Macrophages generate reactive oxygen species in response to minimally oxidized low-density lipoprotein: toll-like receptor 4- and spleen tyrosine kinase-dependent activation of NADPH oxidase 2. Circ Res 104, 210–218, 221p following 218 (2009).

49. J. R. Cubillos-Ruiz et al., ER Stress Sensor XBP1 Controls Anti-tumor Immunity by Disrupting Dendritic Cell Homeostasis. Cell 161, 1527–1538 (2015).

50. E. Vladykovskaya et al., Lipid peroxidation product 4-hydroxy-trans-2-nonenal causes endothelial activation by inducing endoplasmic reticulum stress. J Biol Chem 287, 11398–11409 (2012).

51. B. Kalyanaraman et al., Measuring reactive oxygen and nitrogen species with fluorescent probes: challenges and limitations. Free radical biology & medicine 52, 1–6 (2012).

52. C. Riganti et al., Diphenyleneiodonium inhibits the cell redox metabolism and induces oxidative stress. J Biol Chem 279, 47726–47731 (2004).

53. K. Wingler et al., VAS2870 is a pan-NADPH oxidase inhibitor. Cell Mol Life Sci 69, 3159–3160 (2012).

54. Q. Hu et al., The mitochondrially targeted antioxidant MitoQ protects the intestinal barrier by ameliorating mitochondrial DNA damage via the Nrf2/ARE signaling pathway. Cell death & disease 9, 403 (2018).

55. E. E. To et al., Mitochondrial Reactive Oxygen Species Contribute to Pathological Inflammation During Influenza A Virus Infection in Mice. Antioxid Redox Signal 32, 929–942 (2020).

56. J. de Boer et al., Transgenic mice with hematopoietic and lymphoid specific expression of Cre. Eur J Immunol 33, 314–325 (2003).

57. S. E. Bettigole et al., The transcription factor XBP1 is selectively required for eosinophil differentiation. Nat Immunol 16, 829–837 (2015).

58. D. M. MacCallum, L. Castillo, A. J. Brown, N. A. Gow, F. C. Odds, Early-expressed chemokines predict kidney immunopathology in experimental disseminated Candida albicans infections. PLoS One 4, e6420 (2009).

59. S. E. Logue et al., Inhibition of IRE1 RNase activity modulates the tumor cell secretome and enhances response to chemotherapy. Nat Commun 9, 3267 (2018).

60. X. Sheng et al., IRE1alpha-XBP1s pathway promotes prostate cancer by activating c-MYC signaling. Nat Commun 10, 323 (2019).

61. N. Zhao et al., Pharmacological targeting of MYC-regulated IRE1/XBP1 pathway suppresses MYC-driven breast cancer. J Clin Invest 128, 1283–1299 (2018).

62. R. Matono, K. Miyano, T. Kiyohara, H. Sumimoto, Arachidonic acid induces direct interaction of the p67(phox)-Rac complex with the phagocyte oxidase Nox2, leading to superoxide production. J Biol Chem 289, 24874–24884 (2014).

63. P. V. Seegren et al., Mitochondrial Ca(2+) Signaling Is an Electrometabolic Switch to Fuel Phagosome Killing. Cell reports 33, 108411 (2020).

64. C. Mancebo et al., Fungal Patterns Induce Cytokine Expression through Fluxes of Metabolic Intermediates That Support Glycolysis and Oxidative Phosphorylation. J Immunol 208, 2779–2794 (2022).

65. M. L. Gil, D. Gozalbo, Role of Toll-like receptors in systemic Candida albicans infections. Front Biosci (Landmark Ed) 14, 570–582 (2009).

66. M. G. Netea et al., Toll-like receptor 2 suppresses immunity against Candida albicans through induction of IL-10 and regulatory T cells. J Immunol 172, 3712–3718 (2004).

67. M. J. Elder, S. J. Webster, D. L. Williams, J. S. Gaston, J. C. Goodall, TSLP production by dendritic cells is modulated by IL-1beta and components of the endoplasmic reticulum stress response. Eur J Immunol 46, 455–463 (2016).

68. M. Rodriguez et al., The unfolded protein response and the phosphorylations of activating transcription factor 2 in the trans-activation of il23a promoter produced by beta-glucans. J Biol Chem 289, 22942–22957 (2014).

69. Y. Xiao et al., Targeting CBLB as a potential therapeutic approach for disseminated candidiasis. Nat Med 22, 906–914 (2016).

70. G. Wirnsberger et al., Inhibition of CBLB protects from lethal Candida albicans sepsis. Nat Med 22, 915–923 (2016).

71. X. Zhao et al., JNK1 negatively controls antifungal innate immunity by suppressing CD23 expression. Nat Med 23, 337–346 (2017).

72. A. M. Keestra-Gounder et al., NOD1 and NOD2 signalling links ER stress with inflammation. Nature 532, 394–397 (2016).

73. G. Sule et al., Endoplasmic reticulum stress sensor IRE1alpha propels neutrophil hyperactivity in lupus. J Clin Invest 131, (2021).

74. G. D. Brown et al., Hidden killers: human fungal infections. Sci Transl Med 4, 165rv113 (2012).

75. S. V. Tsay et al., Burden of Candidemia in the United States, 2017. Clin Infect Dis 71, e449–e453 (2020).

76. H. R. Conti, A. R. Huppler, N. Whibley, S. L. Gaffen, Animal models for candidiasis. Curr Protoc Immunol 105, 19 16 11–17 (2014).

77. T. Iwawaki, R. Akai, S. Yamanaka, K. Kohno, Function of IRE1 alpha in the placenta is essential for placental development and embryonic viability. Proc Natl Acad Sci U S A 106, 16657–16662 (2009).

78. S. Saijo et al., Dectin-1 is required for host defense against Pneumocystis carinii but not against Candida albicans. Nat Immunol 8, 39–46 (2007).

79. N. N. Iwakoshi et al., Plasma cell differentiation and the unfolded protein response intersect at the transcription factor XBP-1. Nat Immunol 4, 321–329 (2003).

80. A. M. Gillum, E. Y. Tsay, D. R. Kirsch, Isolation of the Candida albicans gene for orotidine-5’-phosphate decarboxylase by complementation of S. cerevisiae ura3 and E. coli pyrF mutations. Mol Gen Genet 198, 179–182 (1984).

81. M. Behnen et al., Immobilized immune complexes induce neutrophil extracellular trap release by human neutrophil granulocytes via FcgammaRIIIB and Mac-1. J Immunol 193, 1954–1965 (2014).

82. D. Awasthi et al., Glycolysis dependent lactate formation in neutrophils: A metabolic link between NOX-dependent and independent NETosis. Biochim Biophys Acta Mol Basis Dis 1865, 165542 (2019).

83. M. B. Greenblatt, A. Aliprantis, B. Hu, L. H. Glimcher, Calcineurin regulates innate antifungal immunity in neutrophils. J Exp Med 207, 923–931 (2010).

84. N. L. Bray, H. Pimentel, P. Melsted, L. Pachter, Near-optimal probabilistic RNA-seq quantification. Nat Biotechnol 34, 525–527 (2016).

85. M. I. Love, W. Huber, S. Anders, Moderated estimation of fold change and dispersion for RNA-seq data with DESeq2. Genome Biol 15, 550 (2014).

86. A. Liberzon et al., Molecular signatures database (MSigDB) 3.0. Bioinformatics 27, 1739–1740 (2011).

87. A. Liberzon et al., The Molecular Signatures Database (MSigDB) hallmark gene set collection. Cell Syst 1, 417–425 (2015).

88. A. Subramanian et al., Gene set enrichment analysis: a knowledge-based approach for interpreting genome-wide expression profiles. Proceedings of the National Academy of Sciences of the United States of America 102, 15545–15550 (2005).

